# Finding stable clusterings of single-cell RNA-seq data

**DOI:** 10.1101/2025.09.17.672302

**Authors:** Victor Klebanoff

## Abstract

Run a UMI count matrix through a pipeline to obtain *n* cell clusters. Suppose that counts for an equal number of additional cells from the same experiment become available. Would including them change the result? Form the matrix containing both sets of counts, obtain *n* clusters, restrict this clustering to the initial cells and compare it with the initial clustering. If they are not consistent, conclude that the initial clustering is unstable.

This is unrealistic, but reverse the perspective: given a clustering, process samples of half of the cells. If their clusters are consistent with those of all cells restricted to the samples, conclude that the clustering is stable.

We use divisive hierarchical spectral clustering and define what may be a novel mapping of the dendrogram to nested clusterings.

Counts are transformed to points in low-dimensional Euclidean space. Positive affinities are defined for points that are k-nearest neighbors. The affinity equals the inverse of the distance between points. Ng, Jordan, and Weiss’ algorithm divides the points into two clusters. The normalized cut measures the clusters’ separation. Recursion generates a dendrogram. Set the length of the branch between a node and its daughters to the normalized cut. Nodes’ distances from the root define the mapping to nested clusterings.

Analysis is performed for all cells and for multiple pairs of complementary samples of cells. For a given number of clusters, each sample’s clustering and clusters are compared with those of the full data set (restricted to the sample). This provides measures of the stability of the clustering and its clusters. For three large data sets, this yielded clusterings compatible with published results, though with fewer clusters. Clusterings of two were judged to be stable.

We conclude that it is feasible to identify stable clusterings of as many as 100,000 cells. Future research should explore using differential expression for validation.

## 1 Introduction

Although the stability – or replicability – of clusterings of gene expression data has been of interest for at least twenty-five years, there is evidently no consensus on how to find stable clusterings of single-cell RNA-seq data presented as UMI counts.

Early work using sampling to evaluate the effect of data variation on clusterings includes that of Levine and Domany [1], Ben-Hur et al. [2], Dudoit and Fridlyand [3], Lange et al. [4], and Tibshirani and Walther [5]. Hennig [6] discussed evaluating stability by using samples, as well as by replacing points with noise or by jittering. He observed that “stability … is strongly dependent on the data set. In the same clustering, some clusters may be very stable and others may be extremely unstable.”

More recently, Lun [7] discussed bootstrapping scRNA-seq data, Peyvandipour et al. [8] and Tang et al. [9] studied cluster stability for single-cell data with the objective of identifying (novel) cell types, and Patterson-Cross et al. [10] proposed a framework to evaluate the influence on clustering stability of parameters input to an analytical pipeline. The focus of the paper by Masoero et al. [11] is the replicability of clustering results across studies. It opens with a survey of issues and methods for a single study.

A simple question motivated this work: *If data for twice as many cells were available, would clustering results change?* While this is unknowable, the problem can be approached by reversing the question: *Does using half of the cells give consistent clusterings?* Informally: cluster all of the cells, then randomly divide the set of cells in two, and cluster each subset. Restrict the clustering of all cells to each subset and compare it with the subsets’ clusterings. Repeat for multiple samples. If agreement is good for enough samples, the clustering may be considered stable.

We construct a pipeline that takes a UMI count matrix as input and produces clusterings of a range of *sizes* (number of clusters). Next, clusterings are generated for several samples of cells. Comparing the clusterings of the samples with the same-size clusterings of the full set of cells gives stability estimates. Clusterings are compared using what Meilă calls the *misclassification error distance* (MED) [12]. Von Luxburg calls it the *minimal matching distance* [13]. Its distribution across samples characterizes a clustering’s stability. For each *cluster* and *sample* a membership-based *cluster misclassification error rate* (CMER) is defined. Its distribution across samples characterizes the cluster’s stability.

We defined, but do not use here, a *misclassification error rate for each cell*. In some cases, exploratory analysis found recalcitrant cells – cells that were often misclassified in clusterings of different sizes. Although we did not find an efficient way to identify these before clustering, the idea may merit further study as a way to identify outlier cells.

For gene expression data, differential expression analysis may provide independent evidence of stability. If genes’ expression levels between pairs of clusters found to be stable by our method differ consistently across samples, it could corroborate our results. Comparing samples may exempt this from Zhang’s critique [14] of performing clustering and differential expression analysis with the same data.

## 2 Methods

### 2.1 Overview

Suppose that the UMI count matrix ***U*** for a set of cells ***C*** is run through a pipeline to produce a clustering of size *n* (alternatively, an *n*-*clustering*). Prepare two *complementary samples* of cells: randomly assign half to the subset ***C***_**1**_ and the balance to ***C***_**2**_. Prepare the sub-matrix ***U***_**1**_ with counts for cells in ***C***_**1**_. Run ***U***_**1**_ through the pipeline to obtain an *n*-clustering. If the clustering of ***C*** restricted to ***C***_**1**_ is close to the clustering of ***C***_**1**_ alone, we might conclude that *if data for twice as many cells were available* – i.e. combining ***C***_**1**_ and ***C***_**2**_ – *the clustering results would not change*. This answers our motivating question for ***C***_**1**_.

Multiple samples ***C***_***s***_ are clustered using the counts ***U***_***s***_. The MED between the clustering of ***C***_***s***_ and the restriction of the clustering of ***C*** to ***C***_***s***_ is calculated for each *s*. MED is normalized to adjust for the chance grouping of cells. This is analogous to calculating the adjusted Rand index. If the values of MED are sufficiently small with enough samples, consider the clustering stable. In this article, we propose calling a clustering stable if the 90^*th*^ percentile of normalized MED is less than or equal to 0.10. Continue by calculating each cluster’s CMER with all samples. Like MED, CMER is normalized. If the values of CMER are sufficiently small with enough samples, consider the cluster stable. We propose calling a cluster stable if the 90^*th*^ percentile of normalized CMER is less than or equal to 0.50: with at least 90% of the samples, fewer than half of the cells in the cluster are misclassified.

Because – as noted by Hennig (quoted in our Introduction) – a clustering may include clusters that are very stable and others that are extremely unstable, it is necessary to decide when unstable clusters render a clustering unsuitable for downstream analysis. We arbitrarily consider a stable clustering *admissible* for downstream analysis if its unstable clusters have fewer than 500 cells.

If more than one admissible clustering is found for a data set, we generally review results for the one with the most clusters.

### 2.2 Data sets

Seven publicly available data sets were studied (Table 1). The Zhengmixeq data [15] were prepared for a study that evaluated clustering algorithms’ ability of recovering known cell subpopulations [16]. The PBMC data were studied by Zheng et al. in their seminal paper [17]. The other four data sets have recently been used to evaluate proposed analytical methods. Lause et al. [18] studied the retinal data [19]. Grabski et al. [20] studied the lung data [21]. Nicol and Miller [22] studied the breast cancer data [23]. They analyzed the monocytes in a preliminary version of their paper [24].

**Table 1.** Data sets used.

| Data set | source | cells | genes |
| --- | --- | --- | --- |
| Zhengmix4eq | Bioconductor | 3,994 | 15,568 |
| Zhengmix8eq | (same) | 3,994 | 15,716 |
| 68k PBMC | 10x Genomics | 68,579 | 32,738 |
| CD14 Monocytes | 10x Genomics | 2,612 | 32,738 |
| 25k retinal | GSE63472 | 24,769 | 22,292 |
| 65k lung | synapse.org | 65,662 | 26,485 |
| 100k breast cancer | GSE176078 | 100,064 | 29,733 |

Batch correction, at the patient level, was performed for the lung and breast cancer data.

### 2.3 Filter and transform counts; reduce dimensionality

The first step in the pipeline is preliminary filtering. We follow the model of the Seurat Guided Clustering Tutorial [25] – restricting to genes with nonzero counts on at least 50 cells and, for two of the three data sets that were input through Seurat [26] (PBMC and monocytes), excluding cells with high count contributions from mitochondrial genes.

Samples of cells ***C***_***s***_ and the associated count matrices ***U***_***s***_ are prepared. Twenty pairs of complementary samples are used, 40 samples in all, to strike a balance between computing time and having sufficient data for meaningful results.

Following the suggestion in [27] that “applying PCA to deviance or Pearson residuals provides a useful and fast approximation to GLM-PCA,” UMI counts are normalized as Pearson residuals under a Poisson model.

The variability of each gene *g* is calculated with all counts ***U*** – and with each sample’s counts ***U***_***s***_ – as the *sum of squares (SSQ) of its Pearson residuals (PR)*, denoted *S*_*g*_ for the full set and *S*_*g*_(*s*) for sample *s*. These computations are efficiently performed with sparse arrays. The Pearson residuals matrices are not required. The calculation applies a formula derived in Appendix A of our 2023 bioRxiv posting [28] which is included for convenience as Appendix A of this article.

Genes are retained for downstream analysis if they are highly variable – arbitrarily defined as being among the top 2,000 – for the full set of cells and every sample. For the data sets studied, this retained between 565 and 1,704 genes, referred to as *analysis genes*.

To compare genes’ variability across samples or between data sets, it is convenient to normalize *S*_*g*_ by dividing by the number of cells in the count matrix. This yields the mean SSQ of PR, denoted *M*_*g*_.

The Pearson residuals matrix is explicitly calculated for the analysis genes. It is viewed as a low rank matrix perturbed by noise. To estimate the rank of the unperturbed matrix, Erichson’s optht program [29] is used. It implements Gavish and Donoho’s algorithm [30]. For the data sets studied, the ranks of the Pearson residuals range between 11 and 434. The singular value decomposition produces the low rank *SVD representation* of the residuals. By *the distance between cells* we mean the distance between the corresponding points in the SVD representation. By the *diameter* of the set of cells we mean the maximum distance between points in the SVD representation.

### 2.4 Exclude k-nearest neighbor outliers; cluster cells

To cluster points in the SVD representation, the Leiden algorithm was initially considered due to its use in Seurat. This presents difficulties because of the need to vary the resolution parameter to obtain all clusterings of a range of sizes. Suppose, for simplicity, that resolution equal to 2 gives a 2-clustering and resolution equal to 10 gives a 10-clustering. Resolution values between 2 and 10 must be evaluated systematically to obtain all clusterings of sizes 3 through 9. In some cases, the number of clusters failed to increase monotonically with resolution. As a result, some clusterings were not found for some samples.

We explored spectral clustering and developed a divisive hierarchical scheme. Although an exhaustive study was not performed, it appears that the hierarchical approach is more likely to deliver stable clusterings than an alternative that directly outputs clusterings of each size.

Ng, Jordan, and Weiss’ [31] algorithm is used, with a modification: the affinity between two points is defined not by a Gaussian function of the distance between them but as the inverse of the distance. (In section 2.8 of [32], the distance between points is defined as the inverse of their affinity.) Nonzero affinities are defined only for points that are k-nearest neighbors (kNN). In the language of section 14.5.3 *Spectral Clustering* of [33], the *mutual K-nearest-neighbor graph* describes the linkages between points. The affinity matrix is sparse – keeping computations affordable – at the cost of arbitrarily specifying the number of neighbors to be included. While we cannot offer a principled basis for specifying a value of this parameter, restricting to 64^*th*^-nearest neighbors gave credible results. Extensive comparisons were not performed but, in a few cases, using 256^*th*^-nearest neighbors did not improve results.

Meilă recommends that outliers be removed before performing spectral clustering [34]. This led us to consider the distribution of Euclidean distance between points in the SVD representation that are kNN.

These results may have implications for other clustering schemes. Illustrating with the PBMC data (details are in section 3.4), the Euclidean distance between nearest neighbors ranges from 1.3 to 294 (Table 5). If an algorithm relies only on the kNN graph – otherwise ignoring distance – two nearest neighbors 200 units apart are treated no differently than a pair separated by 2 units. To address this, if two points are kNN for specified values of *k*, the affinity is defined as the inverse of the distance between them. For points 200 units apart, the affinity is 0.005. For points two units apart, it equals 0.5.

Returning to Meilă’s recommendation, we propose excluding not only nearest neighbor outliers, but also small sets of isolated points. For example, consider a set of points {*P*_1_, …, *P*_33_} representing 33 cells and suppose that the distance between each pair of points is less than one, but that their distances to points outside of the set are much larger. The 32^*nd*^-nearest neighbor of each *P*_*i*_ belongs to the set but its 64^*th*^-nearest neighbor does not. The 33 points constitute an isolated set. Its members might be excluded as outliers based on criteria proposed below if the 64-NN distances are sufficiently large. This poses questions that may require further study. Some isolated sets of points may represent cell clusters of biological interest. The challenge is to find a balance between retaining useful data and excluding outliers. Here we may err by excluding too many cells.

To identify outliers, the Euclidean distances between all pairs of points are calculated. Ranking the distances from a point to all others yields the distances to its to k-nearest neighbors. We restrict to *k* = 1, 2, 4, …, *k*_*max*_ – arbitrarily setting *k*_*max*_ = 64. Next, the mean and standard deviation of the distances are calculated for each *k*. The sum of the mean plus three times the standard deviation is the threshold for defining outliers. If the distance from a point to its kNN is larger, the point is excluded as a *kNN outlier*. The points remaining in the SVD representation correspond to cells to be clustered.

Data reduction for each subset ***C***_***s***_ uses results found for the full set of cells. The same analysis genes are used to compute Pearson residuals, which are reduced to the rank calculated by optht. KNN outliers are excluded from each subset.

The affinity matrices for spectral clustering are constructed for the full set of points and for each sample. If two points are 64^*th*^-NN or closer, the affinity is set to the inverse of the Euclidean distance between them. Otherwise the affinity equals zero. Although the property of being kNN is not symmetric (if *P*_2_ is the nearest neighbor of *P*_1_, *P*_1_ may not be the nearest neighbor of *P*_2_), affinity matrices are constructed to be symmetric.

For each set of points to be clustered, divisive hierarchical spectral clustering produces a dendrogram. Its maximum depth and minimum node (cluster) size are specified as *stopping conditions*. For the three small data sets, maximum depth was set to 6 and the minimum cluster size to 50. For the four large data sets, the maximum depth was set to 10. For the retinal data set (the smallest of the four), the minimum cluster size was set to 100; for the others, to 200.

Clustering begins by splitting the set of points into two clusters. Each cluster is split in turn. The process continues until a stopping condition is satisfied: the maximum specified depth is reached or splitting would produce a cluster smaller than the minimum permitted size. This produces a dendrogram for the set of all cells and one for each sample.

### 2.5 Map dendrograms to sets of nested clusterings

This method may be novel.

The *normalized cut*, described by Meilă in [34] and von Luxburg in [35], is used to prioritize the split at a node. The smaller it is, the better the separation between the node’s daughter clusters – and the higher the priority of the split. The length of each branch from a node to its daughters is set to the normalized cut.

After assigning lengths to all branches, each node’s distance to the root is calculated and nodes are sorted by their distance from the root. Map the dendrogram to a set of nested clusterings: the root’s daughters correspond to the 2-clustering. Descend to the pair of nodes next closest to the root. They split the points at their parent (one of the root’s daughters), yielding the 3-clustering. Repeat by descending the dendrogram to the pair of nodes next closest to the root – and use them to obtain the 4-clustering. Continue until the maximum specified number of clusters have been found, or the clustering consisting of all terminal nodes is defined.

### 2.6 Calculate quantities that characterize stability

Given *N*_*max*_ we want to evaluate the stability of each clustering of size *n* for 2 ≤ *n* ≤ *N*_*max*_. We require clustering results for all 40 samples. The stopping conditions may prevent the generation of a clustering of size *n* for some sample. If this occurs, only smaller clusterings are analyzed. For the three small data sets, *N*_*max*_ was set to 10. For the four large data sets, values larger than the published number of clusters were used: 25 for PBMC, 70 for the others. Given *n*, the clustering of ***C***_***s***_ is compared with the restriction of the clustering of ***C*** to ***C***_***s***_ by computing the misclassification error distance (MED). Following Lange et al. [4] (as pointed out by von Luxburg in [13]) the MED is normalized. It is divided by the mean of the MED calculated with shuffled cluster labels in the samples. As illustrated by the bottom two plots in Figure 2.1 of [13], normalized MED can exceed 1. In this event, it is set to 1 so that our graphical summaries are uniform, as in the top line of Figure 5. Normalized CMER is also capped at 1.

Using MED to compare clusterings assigns labels to the sample’s clusters that are compatible with the cluster labels of the full set of cells. This is key to defining CMER for each cluster *c* and sample *s*:

- Let ***c*** denote the members of cluster *c* in the full set ***C***.
- Let ***c***_***s***_ denote the intersection of ***c*** and ***C***_***s***_.
- Consider the cluster label assigned in ***C***_***s***_ to a cell in ***c***_***s***_.
- If it does not equal *c*, the cell is misclassified in ***C***_***s***_.
- Define the CMER of cluster *c* in sample *s* as the fraction of cells in ***c***_***s***_ that are misclassified.

Repeat with shuffled cluster labels to calculate denominators for normalizing CMER.

The confusion matrix displayed in Table 2 illustrates the calculation of *unnormalized* MED and CMER for a clustering of the PBMC data that is reviewed in section 3.5. (This is the 9-clustering produced in the third iteration.)

**Table 2.** 68k PBMC data (iteration 3): compare the 9-clusterings calculated with all cells and with sample 1.

|  | 0 | 1 | 2 | 3 | 4 | 5 | 6 | 7 | 8 | Total |
| --- | --- | --- | --- | --- | --- | --- | --- | --- | --- | --- |
| 0 | 9,057 | 150 | 0 | 0 | 0 | 71 | 0 | 0 | 0 | 9,278 |
| 1 | 32 | 2,907 | 0 | 0 | 0 | 3 | 0 | 0 | 0 | 2,942 |
| 2 | 0 | 0 | 1,825 | 0 | 4 | 0 | 5 | 0 | 0 | 1,834 |
| 3 | 0 | 0 | 145 | 0 | 9 | 0 | 0 | 0 | 0 | 154 |
| 4 | 0 | 0 | 4 | 0 | 2,618 | 0 | 27 | 0 | 0 | 2,649 |
| 5 | 74 | 4 | 0 | 0 | 0 | 8,383 | 5 | 0 | 0 | 8,466 |
| 6 | 0 | 1 | 1 | 0 | 11 | 146 | 2,779 | 0 | 0 | 2,938 |
| 7 | 1 | 0 | 0 | 0 | 0 | 1 | 0 | 1,440 | 0 | 1,442 |
| 8 | 0 | 0 | 0 | 211 | 0 | 1 | 2 | 0 | 1,598 | 1,812 |
| Total | 9,164 | 3,062 | 1,975 | 211 | 2,642 | 8,605 | 2,818 | 1,440 | 1,598 | 31,515 |

The table compares the cluster assignments calculated with all cells (row labels) with the assignments calculated with a sample containing 31,515 cells (column labels).

Columns are sorted to maximize the sum of the diagonal. *This assigns the cluster labels in the sample*.

- The MED is the fraction of the total count that is off the diagonal, here equal to 0.029.
- In the fourth row – counts in the sample for cells assigned to cluster 3 in the full set of data – the diagonal entry equals zero. All 154 of these cells are misclassified in the sample; CMER=1.
- In the second row – counts in the sample for cells assigned to cluster 1 in the full set of data – the diagonal entry equals 2,907. 35 cells are misclassified in the sample; CMER=0.01.
- In the seventh row – counts in the sample for cells assigned to cluster 6 in the full set of data – the diagonal entry equals 2,779. 159 cells are misclassified in the sample; CMER=0.05.
- In the bottom row – counts in the sample for cells assigned to cluster 8 in the full set of data – the diagonal entry equals 1,598. 214 cells are misclassified in the sample; CMER=0.12.

Compare with the confusion matrix in Table 3 – for a sample containing 31,552 cells:

- MED equals 0.178 – six times larger.
- Cluster 3: all 133 cells are misclassified; CMER=1; identical to Table 2.
- Cluster 1: all 2,934 cells are misclassified, making CMER=1; in Table 2, CMER=0.01.
- Cluster 6: CMER=0.46; in Table 2, CMER=0.05.
- Cluster 8: CMER=0.44; in Table 2, CMER=0.12.

**Table 3.** 68k PBMC data (iteration 3): compare the 9-clusterings calculated with all cells and with sample 32.

|  | 0 | 1 | 2 | 3 | 4 | 5 | 6 | 7 | 8 | Total |
| --- | --- | --- | --- | --- | --- | --- | --- | --- | --- | --- |
| 0 | 9,195 | 0 | 0 | 0 | 0 | 183 | 0 | 0 | 0 | 9,378 |
| 1 | 2,931 | 0 | 0 | 0 | 0 | 3 | 0 | 0 | 0 | 2,934 |
| 2 | 0 | 0 | 1,809 | 0 | 11 | 0 | 1 | 0 | 0 | 1,821 |
| 3 | 0 | 0 | 128 | 0 | 5 | 0 | 0 | 0 | 0 | 133 |
| 4 | 0 | 0 | 5 | 0 | 2,617 | 0 | 1 | 0 | 0 | 2,623 |
| 5 | 26 | 0 | 0 | 115 | 0 | 8,234 | 21 | 0 | 0 | 8,396 |
| 6 | 0 | 0 | 1 | 1,259 | 118 | 5 | 1,636 | 0 | 0 | 3,019 |
| 7 | 1 | 0 | 1 | 0 | 0 | 1 | 0 | 1,438 | 0 | 1,441 |
| 8 | 0 | 793 | 0 | 0 | 0 | 1 | 1 | 0 | 1,012 | 1,807 |
| Total | 12,153 | 793 | 1,944 | 1,374 | 2,751 | 8,427 | 1,660 | 1,438 | 1,012 | 31,552 |

Motivated by the large CMER values for clusters 3, 1, 6, and 8, we checked the other 38 samples. The following results are for all 40 samples.

- Cluster 3: CMER=1 in 39 samples and 0.9 in one sample. In every sample, cluster 3 is merged with cluster 2.
- Cluster 1: CMER=1 in 13 samples. In each of these, cluster 1 is merged with cluster 0, as in Table 3. In the remaining samples CMER is less than 0.04.
- Cluster 6: CMER is between 0.4 and 0.55 in 19 samples. In 15 of these samples, a large segment of cluster 6 splits to cluster 3, as in Table 3. In the other 4, the split is to cluster 1. In the remaining 21 samples, CMER <0.07.
- Cluster 8: CMER is between 0.1 and 0.5 in 20 samples. In 2 of these samples, a large segment of cluster 8 splits to cluster 1, as in Table 3. In the other 18, the split is to cluster 3, as in Table 2. In the remaining 20 samples, CMER <0.0025.

In [7] Lun proposed comparing pairs of clusters to evaluate pairwise instability – to identify clusters that are *not stable with respect to each other*, that are likely to be merged in a random sample. This is the behavior seen here for the merging of cluster 1 with 0 and of cluster 3 with 2.

A second benefit of assigning cluster labels in the samples that are compatible with the labels in the full set of cells is that it facilitates comparing differential expression across samples. We anticipate that validating our approach will require verifying that the differential expression of analysis genes between pairs of stable clusters is consistent across all samples.

### 2.7 Perform single-factor batch correction, if appropriate

For batch correction, the formula for *S*_*g*_ is modified, corresponding to Nicol and Miller’s calculating a gene-level intercept for each batch ([22] Supplementary material):

- For each batch *b*, independently calculate the SSQ of each gene’s Pearson residuals. Denote this as 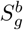 for the set of all cells and as 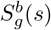 for sample *s*.
- Define 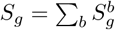 for the full set of cells and 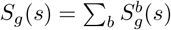 for sample *s*.

Calculate Pearson residuals for the cells in each batch. Concatenate the matrices for downstream analysis.

### 2.8 Identify and exclude cell and gene outliers for iterative analyses

Three sets of analyses were performed for each of the four large data sets. For the first iteration, the count matrices were analyzed as outlined in sections 2.3 through 2.6.

For the second iteration, outliers were excluded from the count matrix. This is motivated by a toy example in our 2023 bioRxiv posting [28] which illustrated the impact of a single outlier cell on the value of *S*_*g*_. In that example, the set of cells is split into two samples. The value of *S*_*g*_ for the subset including the outlier is nine times larger than the value for the complementary sample.

Although we have not explored toy examples for the effect of outliers on clustering, we believe they are impediments to the existence of stable clusterings – that if a count matrix admits a stable clustering, a necessary condition is that the distributions of UMI counts in all sufficiently large samples agree with one another in a sense yet to be discovered. We have not compared the distributions of complementary samples, but have considered properties that a count matrix should have if the distributions of its samples agree. Two, both involving *S*_*g*_(*s*), were used to identify outliers:

1. Cells: If a cell’s contribution to *S*_*g*_(*s*) is large in some sample – i.e. the cell is responsible for an exceptionally large fraction of *S*_*g*_(*s*) – it is an outlier because it cannot have a comparable impact in the complementary sample. The derivation in Appendix B shows how each cell’s fractional contribution to *S*_*g*_ can be calculated for the Poisson model. Each cell’s maximum fractional contribution to *S*_*g*_(*s*) is calculated across all genes and samples.
2. Genes: The toy example suggests that for each gene *g*, the values of *S*_*g*_(*s*) should be consistent across all samples *s*. To evaluate this, calculate the ratio *max*_*s*_*S*_*g*_(*s*)*/min*_*s*_*S*_*g*_(*s*) for each gene. If it is close to one, values of *S*_*g*_(*s*) are consistent. If it is exceptionally large, *g* is an outlier.

After excluding cell and gene outliers, the second iteration followed sections 2.3 through 2.6.

For the third iteration, the counts input to the second iteration were subjected to additional filtering. Cells excluded as kNN outliers during that iteration were removed. This was followed by another round of deleting outlier cells and genes. The resulting counts were analyzed per sections 2.3 through 2.6.

## 3 Results

### 3.1 Impact of data reduction

Table 4 summarizes the impact of initial data reduction. This does not include the effect of excluding outliers as discussed in section 2.8.

**Table 4.** Impact of data reduction.

| Data set | preliminary filtering |  | analysis<br>genes | estimated<br>rank | cells<br>retained |
| --- | --- | --- | --- | --- | --- |
|  | cells | genes |  |  |  |
| Zhengmix4eq | 3,980 | 5,837 | 646 | 35 | 3,902 |
| Zhengmix8eq | 3,974 | 6,075 | 565 | 37 | 3,906 |
| 68k PBMC | 68,286 | 12,515 | 652 | 48 | 67,792 |
| CD14 monocytes | 2,558 | 3,726 | 719 | 11 | 2,537 |
| 25k retinal | 24,769 | 13,552 | 1,081 | 51 | 24,101 |
| 65k lung | 60,993 | 17,470 | 1,659 | 305 | 60,114 |
| 100k breast cancer | 100,064 | 21,354 | 1,704 | 434 | 98,681 |

- Columns 1 and 2: the number of cells and genes retained after preliminary filtering. The Pearson residuals matrix has a column for each cell. Compared to Table 1, the number of genes is smaller because genes with nonzero counts on fewer than 50 cells were excluded. The number of cells is smaller for the Zhengmixeq data because some barcodes have duplicate values (see section 3.5). Cells with high count contributions from mitochondrial genes were excluded from the PBMC and monocyte data. Blood cells were excluded from the lung data.
- Column 3: the number of analysis genes – genes that are highly variable in every sample and the full data set. This is the number of rows of the Pearson residuals matrix.
- Column 4: the rank of the Pearson residuals matrix estimated by the optht program. This is the number of rows in the SVD representation matrix.
- Column 5: the number of cells retained for clustering after excluding kNN outliers.

### 3.2 Genes that are highly variable for all cells may not be highly variable for every sample

The plots in Figure 1 show the relation between the mean SSQ of Pearson residuals calculated with all cells (*M*_*g*_) and the number of cells with nonzero counts for genes retained after preliminary filtering. Black points represent analysis genes retained for downstream analysis: *S*_*g*_ and *S*_*g*_(*s*) are among the 2,000 largest when calculated with all cells and with each sample *s*, respectively.

**Figure 1.**
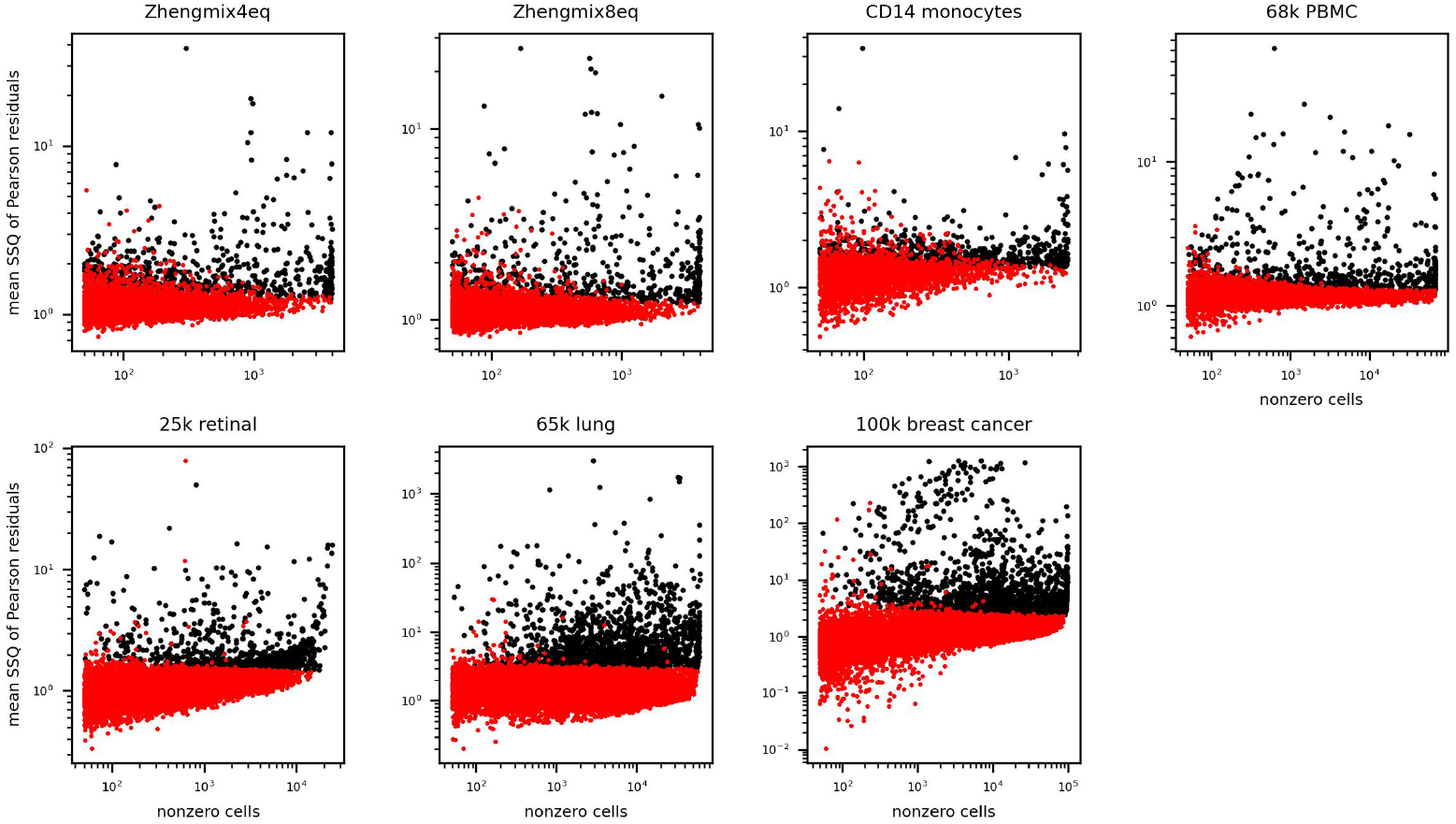
Filtered genes: *M*_*g*_ vs. number of cells with counts *>*0; analysis genes in black

Some genes that are highly variable in the full set of data (large vertical coordinates) are not highly variable in every sample (red points). It is possible that some of these genes would be retained after excluding outlier cells as described in section 2.8.

### 3.3 Analytic estimates of the rank of the Pearson residuals

Ranks of Pearson residuals matrices were estimated with the optht program. Four other programs were considered. Although all five gave reasonable results on toy problems (matrices of known low rank with added noise) each of the others gave problematic results:

- Wide variation between results depending on the user-selected algorithm (one program)
- Rank estimates differing by an order for magnitude for very similar input matrices (one program)
- Long run times (two programs)
- Failure to find a solution (one program)

Results are tabulated in column 4 of Table 4 and plotted in Figure 2 using the format of the screeplot program in the Bioconductor PCAtools package [36]. The estimated ranks of the Pearson residuals for the lung and breast cancer data (305 and 434, respectively) are larger than values we have seen in the literature. The maximum value plotted on the horizontal axis is the number of analysis genes – the number of rows in the Pearson residuals matrix (column 3 of Table 4).

**Figure 2.**
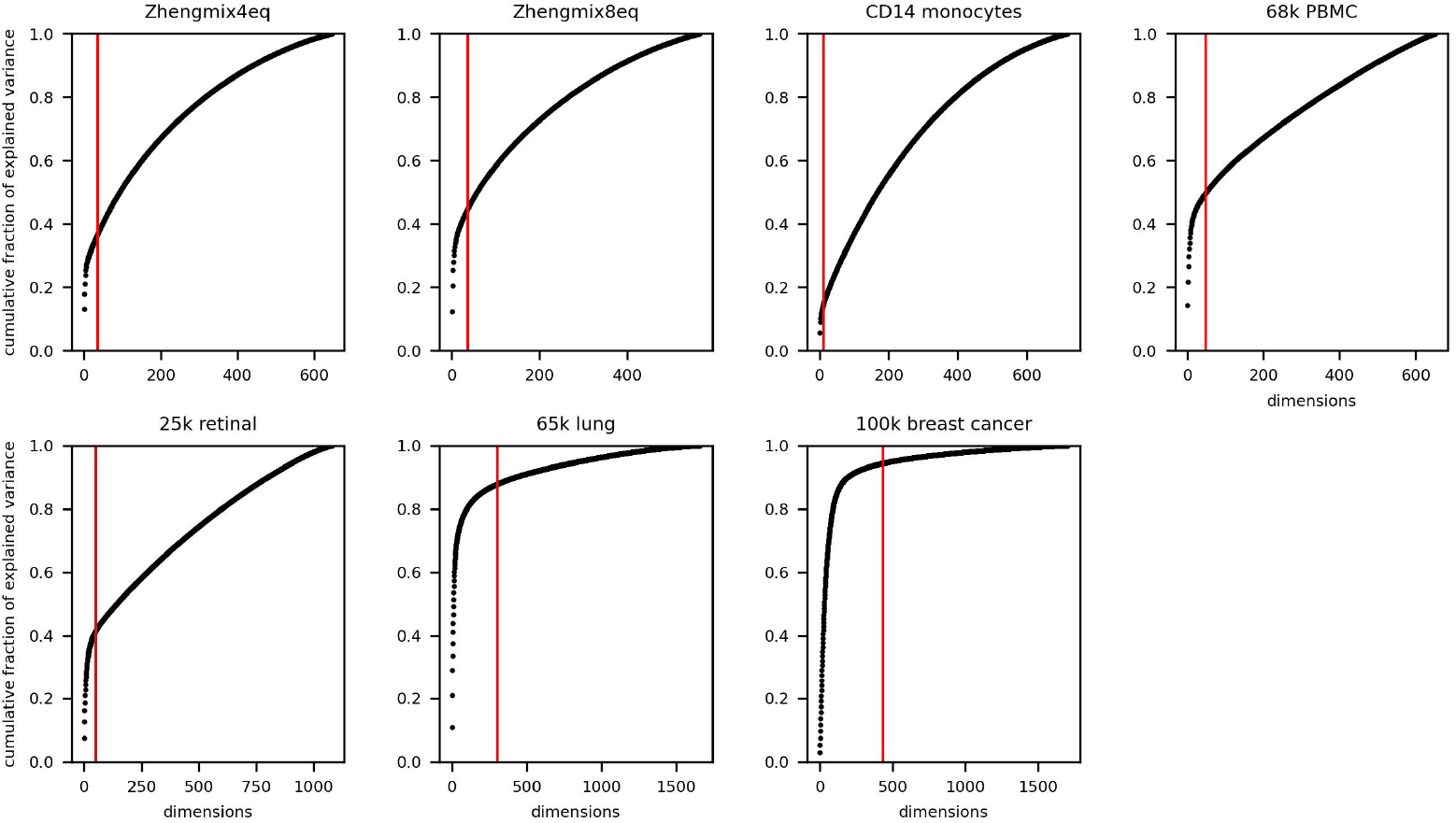
Scree plots: red vertical lines indicate the rank estimated by optht

### 3.4 Variation of Euclidean distance between k-nearest neighbors

Our interest in the relation between kNN and Euclidean distance was motivated by Meilă’s recommendation to exclude outliers before performing spectral clustering and by the work of Cooley et al. [37].

We illustrate with the PBMC and breast cancer data. The variation of Euclidean distance between k-nearest neighbors in the SVD representation of the PBMC data is summarized in Table 5. The first column contains statistics for the distance from a cell to its nearest neighbor, which ranges from 1.3 to 294, with a mean of 5.0 and standard deviation of 7.4. Subsequent columns list statistics for 2^*nd*^-nearest neighbor distance, 4^*th*^-nearest, continuing to 64^*th*^-nearest, and finally the maximum distance between cells. The diameter of the set equals 823 – the bottom right entry.

**Table 5.** 68k PBMC data: distributions of distance between kNN.

|  | 1 | 2 | 4 | 8 | 16 | 32 | 64 | maximum |
| --- | --- | --- | --- | --- | --- | --- | --- | --- |
| mean | 5.0 | 5.4 | 5.8 | 6.1 | 6.6 | 7.1 | 7.7 | 707.9 |
| stddev | 7.4 | 8.0 | 8.7 | 9.7 | 10.7 | 11.8 | 13.4 | 6.3 |
| min | 1.3 | 1.6 | 1.8 | 2.0 | 2.1 | 2.3 | 2.4 | 507.1 |
| 10% | 2.3 | 2.4 | 2.6 | 2.7 | 2.9 | 3.0 | 3.2 | 707.7 |
| 50% | 3.8 | 4.0 | 4.2 | 4.5 | 4.7 | 5.0 | 5.3 | 708.2 |
| 75% | 5.5 | 5.9 | 6.3 | 6.6 | 7.0 | 7.4 | 7.9 | 708.4 |
| 90% | 8.2 | 8.8 | 9.4 | 10.0 | 10.7 | 11.5 | 12.4 | 708.7 |
| 95% | 10.6 | 11.4 | 12.2 | 13.2 | 14.3 | 15.7 | 17.3 | 708.9 |
| 99% | 18.2 | 20.0 | 22.0 | 24.4 | 27.2 | 30.6 | 35.2 | 711.3 |
| max | 294.3 | 384.4 | 389.0 | 433.6 | 461.9 | 507.1 | 551.0 | 822.6 |

Clearly, kNN neighborhoods may not resemble neighborhoods defined with the Euclidean metric. For half of the cells, the distance to the nearest neighbor is less than 4 units – less than 0.5% of the diameter of the set of cells. However, there is a cell whose nearest neighbor is 75 times further away – 294 units distant, 36% of the diameter of the set of cells.

Cells that are exceptionally distant from their kNN are identified as outliers to be excluded. Outliers are defined by distances at least three standard deviations larger than the mean. For nearest neighbors, this threshold equals 27.1. Fewer than 1% of cells are outliers based on this criterion. Applying this to 2^*nd*^, 4^*th*^, …, and 64^*th*^-nearest neighbors excludes a total of 494 cells, retaining 67,792. The distributions of distances for the retained cells are summarized in Table 6. Excluding kNN outliers reduces the range of nearest neighbor distances by an order of magnitude and the diameter of the set by 80%.

**Table 6.**
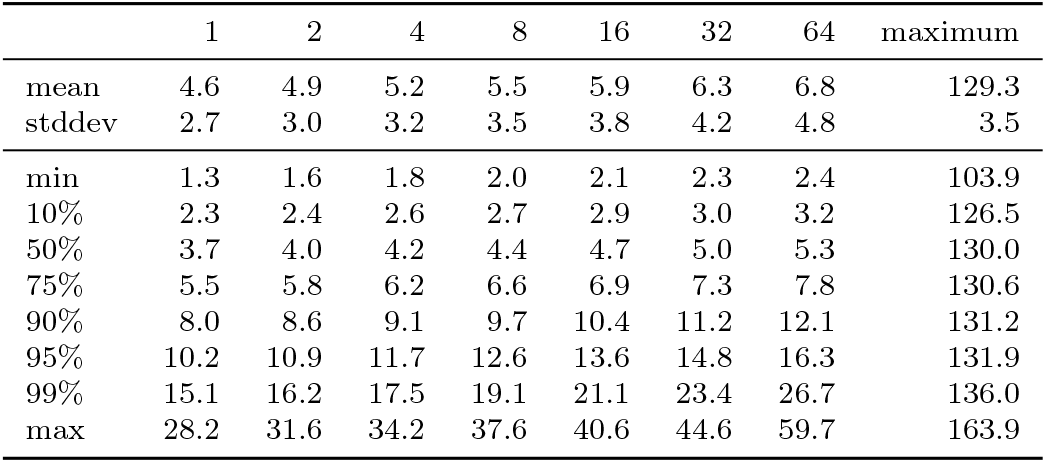
68k PBMC data: distributions of distance between kNN after excluding outliers.

For the breast cancer data, the variation is greater. Before excluding kNN outliers, nearest neighbor distance varies by a factor of 580 (Table 7). Even after excluding 1.4% of the cells, the maximum is 50 times larger than the minimum (Table 8).

**Table 7.**
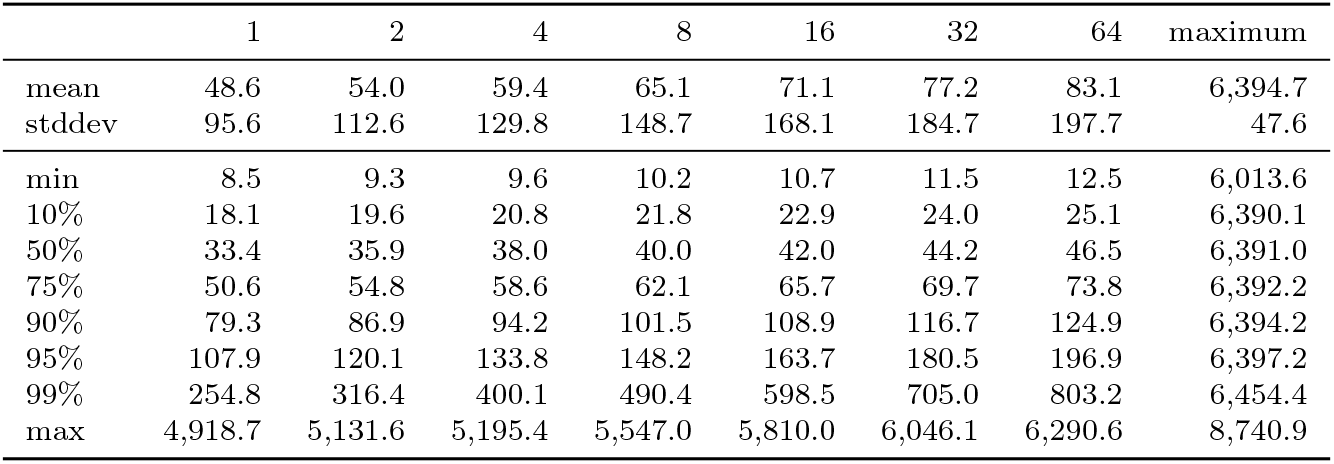
100k breast cancer data: distributions of distance between kNN.

|  | 1 | 2 | 4 | 8 | 16 | 32 | 64 | maximum |
| --- | --- | --- | --- | --- | --- | --- | --- | --- |
| mean | 48.6 | 54.0 | 59.4 | 65.1 | 71.1 | 77.2 | 83.1 | 6,394.7 |
| stddev | 95.6 | 112.6 | 129.8 | 148.7 | 168.1 | 184.7 | 197.7 | 47.6 |
| min | 8.5 | 9.3 | 9.6 | 10.2 | 10.7 | 11.5 | 12.5 | 6,013.6 |
| 10% | 18.1 | 19.6 | 20.8 | 21.8 | 22.9 | 24.0 | 25.1 | 6,390.1 |
| 50% | 33.4 | 35.9 | 38.0 | 40.0 | 42.0 | 44.2 | 46.5 | 6,391.0 |
| 75% | 50.6 | 54.8 | 58.6 | 62.1 | 65.7 | 69.7 | 73.8 | 6,392.2 |
| 90% | 79.3 | 86.9 | 94.2 | 101.5 | 108.9 | 116.7 | 124.9 | 6,394.2 |
| 95% | 107.9 | 120.1 | 133.8 | 148.2 | 163.7 | 180.5 | 196.9 | 6,397.2 |
| 99% | 254.8 | 316.4 | 400.1 | 490.4 | 598.5 | 705.0 | 803.2 | 6,454.4 |
| max | 4,918.7 | 5,131.6 | 5,195.4 | 5,547.0 | 5,810.0 | 6,046.1 | 6,290.6 | 8,740.9 |

**Table 8.** 100k breast cancer data: distributions of distance between kNN after excluding outliers.

|  | 1 | 2 | 4 | 8 | 16 | 32 | 64 | maximum |
| --- | --- | --- | --- | --- | --- | --- | --- | --- |
| mean | 42.2 | 45.8 | 49.2 | 52.8 | 56.6 | 60.7 | 65.1 | 1,189.1 |
| stddev | 31.0 | 34.6 | 38.9 | 44.0 | 50.1 | 57.1 | 65.2 | 18.6 |
| min | 8.5 | 9.3 | 9.6 | 10.2 | 10.7 | 11.5 | 12.5 | 1,050.6 |
| 10% | 18.0 | 19.5 | 20.7 | 21.8 | 22.8 | 23.9 | 25.1 | 1,181.6 |
| 50% | 33.1 | 35.6 | 37.6 | 39.6 | 41.6 | 43.7 | 46.0 | 1,185.3 |
| 75% | 49.6 | 53.5 | 57.1 | 60.6 | 64.1 | 67.9 | 71.9 | 1,189.8 |
| 90% | 75.7 | 82.3 | 89.0 | 95.6 | 102.3 | 109.4 | 116.8 | 1,198.2 |
| 95% | 98.4 | 108.4 | 118.4 | 129.2 | 141.3 | 153.6 | 166.2 | 1,207.7 |
| 99% | 168.5 | 190.1 | 215.0 | 246.6 | 285.3 | 328.3 | 385.9 | 1,266.6 |
| max | 427.9 | 429.6 | 502.6 | 535.2 | 604.7 | 640.8 | 892.8 | 1,566.0 |

### 3.5 Clustering results for individual data sets

In section 2.1 we proposed considering a clustering stable if the 90^*th*^ percentile of normalized MED is less than or equal to 0.10. A cluster is judged stable if the 90^*th*^ percentile of normalized CMER is less than or equal to 0.50. A stable clustering is admissible for downstream analysis if its unstable clusters have fewer than 500 cells.

Begin with the three small data sets. For the Zhengmixeq data we are interested in (1) the relation between the ground truth labels and our method’s clusterings and (2) the stability of specific clusterings and clusters. For the Zhengmix4eq data, agreement with ground truth labels is excellent; for the Zhengmix8eq data less so, though typical of what we have found in publications. For the monocytes, our results suggest that there are no stable clusterings.

For each of the four large data sets, as outlined in section 2.8, three sets of analyses were performed, progressively excluding outlier cells and genes. Six clusterings are reviewed:

- PBMC: two clusterings; an admissible 12-clustering and an unstable 9-clustering
- retinal: an admissible 11-clustering
- lung: two admissible clusterings; one with 19 clusters, the other with 16
- breast cancer: an inadmissible clustering compatible with published results

**Table 9.** Zhengmix4eq data: compare ground truth with the 4-clustering.

| ground truth | hierarchical cluster |  |  |  | Total |
| --- | --- | --- | --- | --- | --- |
|  | 0 | 1 | 2 | 3 |  |
| naive.cytotoxic | 977 | 16 | 0 | 0 | 993 |
| regulatory.t | 5 | 986 | 0 | 0 | 991 |
| b.cells | 0 | 0 | 997 | 0 | 997 |
| cd14.monocytes | 0 | 9 | 2 | 910 | 921 |
| Total | 982 | 1,011 | 999 | 910 | 3,902 |

#### Zhengmix4eq

The data set contains counts for 15,568 genes and 3,994 cells. They represent four cell types. Ground truth labels are provided with the data.

Seven barcodes appear twice. The corresponding 14 columns were dropped, retaining counts for 3,980 cells. Two EnsemblIDs have the same gene symbol (SRSF10). Data for the EnsemblID with nonzero counts on fewer cells were excluded.

Filtering to exclude genes with nonzero counts on fewer than 50 cells retained 5,837 genes. Because data were not input through Seurat, cells were not screened for high mitochondrial DNA levels. 646 genes were found to be highly variable in the full data set and in all 40 samples.

The rank of the Pearson residuals matrix was estimated as 35 by optht. After mapping to a 35-dimensional SVD representation and excluding kNN outliers, 3,902 cells were retained.

Figure 3 displays the distributions of normalized MED for the clusterings of sizes 2-10. There is one line per clustering. The 40 vertically jittered dots in each line show the samples’ MED. For each clustering, the blue vertical segment marks the median of the distribution. The 75^*th*^ percentile marker is green. The 90^*th*^ percentile marker is red. The clusterings of sizes 2-5 are stable. Their 90^*th*^ percentiles are less than or equal to 0.10. For the clusterings of sizes 2-4, the plotted percentiles equal 0.00. Only the blue markers are visible because we favor the lower percentile markers.

**Figure 3.**
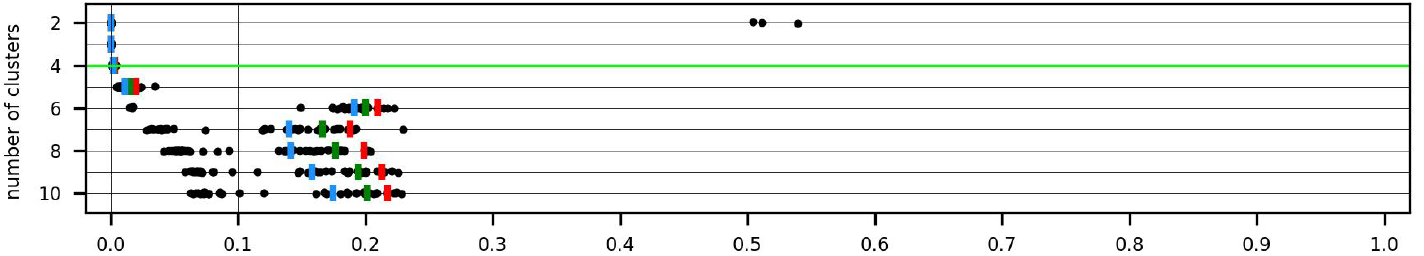
Zhengmix4eq data: distributions of normalized MED for clusterings of sizes 2-10

Because there are 4 ground truth labels, we review the 4-clustering. The green highlighted line displays its MED values.

Figure 4 displays the distributions of normalized CMER for each cluster. All four are stable. The largest value of CMER equals 0.017. The plot shows the median, 75^*th*^, and 90^*th*^ percentiles of CMER for each cluster. They are indistinguishable for clusters 0, 2, and 3. In addition, the colored vertical lines that extend from the bottom to the top of the plot (all very close to 0 on the horizontal axis) indicate the clustering’s percentiles – the ones marked in Figure 3.

**Figure 4.**
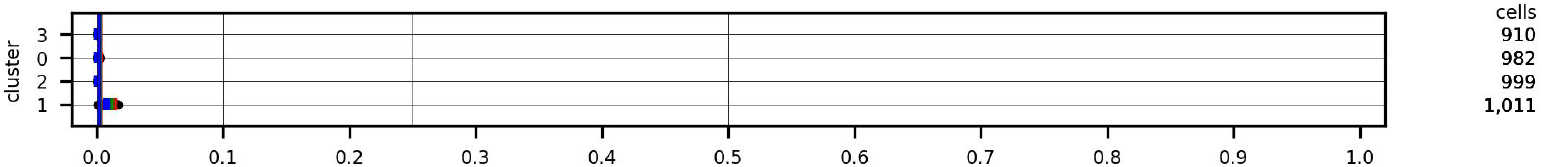
Zhengmix4eq data: distributions of normalized CMER for the 4-clustering

Table 9 compares the clusters with the ground truth labels.

#### Zhengmix8eq

The data set contains counts for 15,716 genes and 3,994 cells. They represent eight cell types or subtypes. Ground truth labels are provided with the data.

Ten barcodes appear twice. The corresponding 20 columns were dropped, retaining counts for 3,974 cells. Two EnsemblIDs have the same gene symbol (SRSF10). Data for the EnsemblID with nonzero counts on fewer cells were excluded.

Filtering to exclude genes with nonzero counts on fewer than 50 cells retained 6,075 genes. Because data were not input through Seurat, cells were not screened for high mitochondrial DNA levels. 565 genes were found to be highly variable in the full data set and in all 40 samples.

The rank of the Pearson residuals matrix was estimated as 37 by optht. After mapping to a 37-dimensional SVD representation and excluding kNN outliers, 3,906 cells were retained.

Figure 5 displays the distributions of normalized MED for the clusterings of sizes 2-10. Differences with Figure 3 are immediate: values for the clusterings of sizes 2 and 3 are large. The clusterings of sizes 4-8 are stable. The clusterings of sizes 7 and 8 are reviewed (green lines).

**Figure 5.**
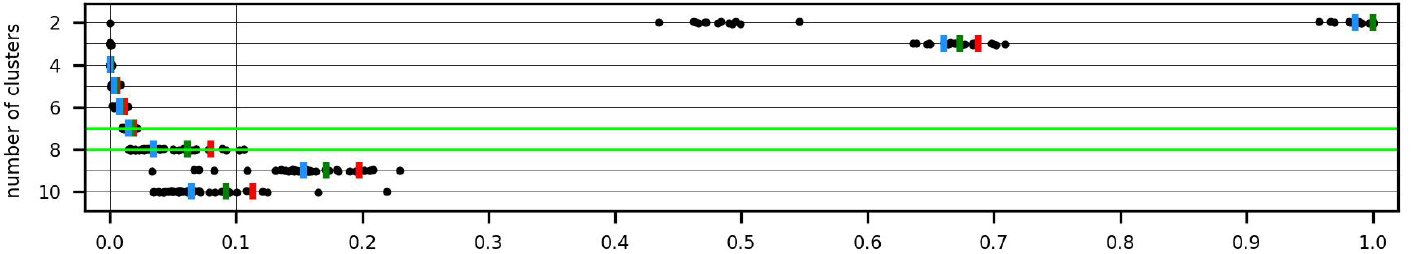
Zhengmix8eq data: distributions of normalized MED for clusterings of sizes 2-10

Figure 6 displays the distributions of normalized CMER for the 7 clusters. All are stable.

**Figure 6.**
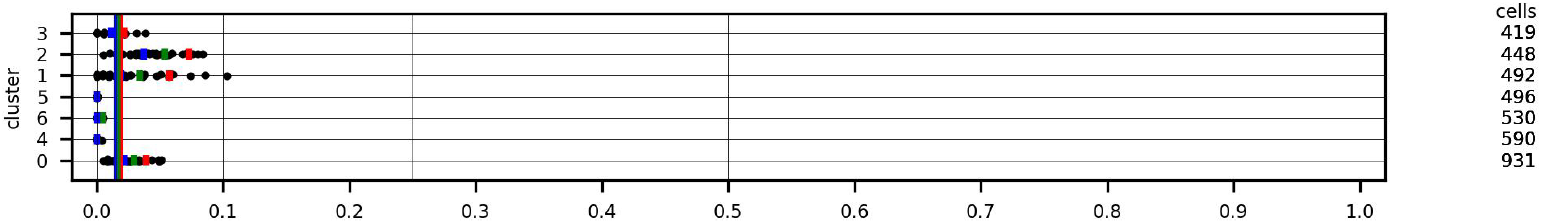
Zhengmix8eq data: distributions of normalized CMER for the 7-clustering

Table 10 compares the clusters with the ground truth labels. Results are very accurate for cd56.nk, b.cells, and cd14.monocytes; less accurate for memory.t and naive.cytotoxic cells. Three subtypes – cd4.t.helper, naive.t, and regulatory.t – are commingled in clusters 0 and 1. The adjusted Rand index equals 0.74.

**Table 10.** Zhengmix8eq data: compare ground truth with the 7-clustering.

| ground truth | hierarchical cluster |  |  |  |  |  |  | Total |
| --- | --- | --- | --- | --- | --- | --- | --- | --- |
|  | 0 | 1 | 2 | 3 | 4 | 5 | 6 |  |
| cd4.t.helper | 258 | 138 | 2 | 0 | 0 | 0 | 0 | 398 |
| naive.t | 485 | 7 | 0 | 2 | 0 | 0 | 0 | 494 |
| regulatory.t | 186 | 308 | 1 | 0 | 0 | 0 | 0 | 495 |
| memory.t | 1 | 33 | 435 | 26 | 0 | 0 | 0 | 495 |
| naive.cytotoxic | 1 | 1 | 5 | 389 | 0 | 1 | 0 | 397 |
| cd56.nk | 0 | 0 | 5 | 2 | 589 | 1 | 0 | 597 |
| b.cells | 0 | 0 | 0 | 0 | 0 | 493 | 0 | 493 |
| cd14.monocytes | 0 | 5 | 0 | 0 | 1 | 1 | 530 | 537 |
| Total | 931 | 492 | 448 | 419 | 590 | 496 | 530 | 3,906 |

The 8-clustering is formed from the 7-clustering by splitting its cluster 4 (590 cells) into the two clusters 0 (339 cells) and 1 (251 cells). Figure 7 displays the distributions of normalized CMER for the 8 clusters. All are stable, but the two new clusters (top two lines) are much less stable than the cluster from which they were formed.

**Figure 7.**
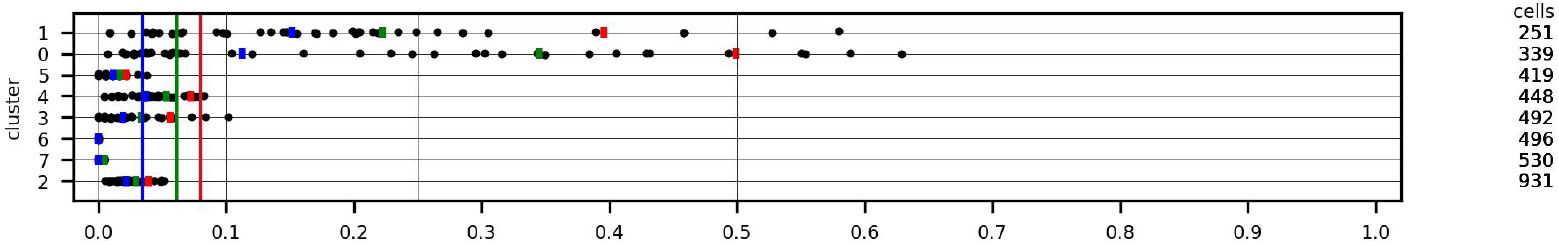
Zhengmix8eq data: distributions of normalized CMER for the 8-clustering

Table 11 compares the clusters with the ground truth labels. The adjusted Rand index equals 0.68.

**Table 11.** Zhengmix8eq data: compare ground truth with the 8-clustering.

| ground truth | hierarchical cluster |  |  |  |  |  |  |  | Total |
| --- | --- | --- | --- | --- | --- | --- | --- | --- | --- |
|  | 0 | 1 | 2 | 3 | 4 | 5 | 6 | 7 |  |
| cd56.nk | 339 | 250 | 0 | 0 | 5 | 2 | 1 | 0 | 597 |
| cd4.t.helper | 0 | 0 | 258 | 138 | 2 | 0 | 0 | 0 | 398 |
| naive.t | 0 | 0 | 485 | 7 | 0 | 2 | 0 | 0 | 494 |
| regulatory.t | 0 | 0 | 186 | 308 | 1 | 0 | 0 | 0 | 495 |
| memory.t | 0 | 0 | 1 | 33 | 435 | 26 | 0 | 0 | 495 |
| naive.cytotoxic | 0 | 0 | 1 | 1 | 5 | 389 | 1 | 0 | 397 |
| b.cells | 0 | 0 | 0 | 0 | 0 | 0 | 493 | 0 | 493 |
| cd14.monocytes | 0 | 1 | 0 | 5 | 0 | 0 | 1 | 530 | 537 |
| Total | 339 | 251 | 931 | 492 | 448 | 419 | 496 | 530 | 3,906 |

#### CD14 Monocytes

The data set contains counts for 32,738 genes and 2,612 cells. Filtering to exclude genes with nonzero counts on fewer than 50 cells and to exclude cells with more than 5% of counts due to mitochondrial DNA retained 3,726 genes and 2,558 cells. 719 genes were found to be highly variable in the full data set and in all 40 samples.

The rank of the Pearson residuals matrix was estimated as 11 by optht. After mapping to an 11-dimensional SVD representation and excluding kNN outliers, 2,537 cells were retained.

Figure 8 displays the distributions of normalized MED for the clusterings of sizes 2-10. No other data set reviewed in this article has so many large values for small clusterings. The median is greater than 0.50 for all clusterings. This is consistent with all cells being of the same type, so that any clustering would be spurious.

**Figure 8.**
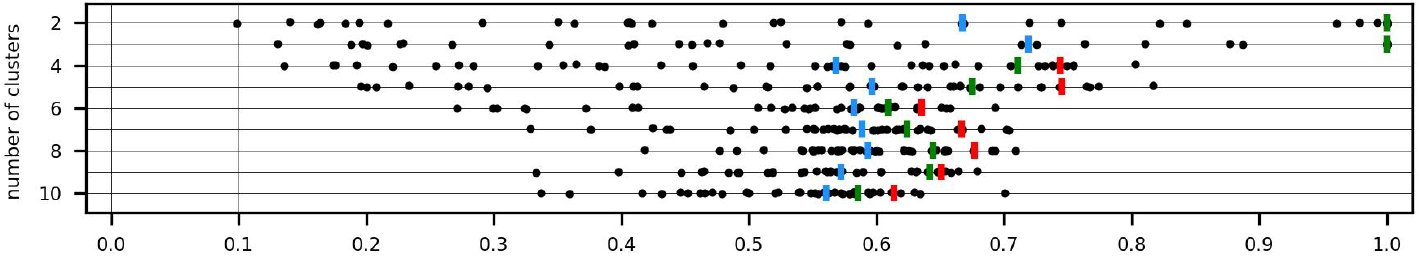
CD14 Monocytes: distributions of normalized MED for clusterings of sizes 2-10

#### 68k PBMC

The data set contains counts for 32,738 genes and 68,579 cells. Filtering to exclude genes with nonzero counts on fewer than 50 cells and to exclude cells with more than 5% of counts due to mitochondrial DNA retained 12,515 genes and 68,286 cells. 652 genes were found to be highly variable in the full data set and in all 40 samples.

The rank of the Pearson residuals matrix was estimated as 48 by optht. After mapping to a 48-dimensional SVD representation and excluding kNN outliers, 67,792 cells were retained.

Our objective was to evaluate clusterings of sizes up to 25 – larger than the 10 reported in the paper [17]. Clusterings of all sizes in the range 2-25 were found in each of the three sets of analyses. For the first iteration, the clusterings of sizes 2, 3, 6, 11, and 12 are stable. The largest is admissible, as shown below. The second iteration, using fewer cells, also yields a stable 12-clustering, but it is not admissible. It has an unstable cluster of 1,716 cells. The third iteration yields two stable clusterings, but they are small, with 2 and 3 clusters.

Figure 9 displays the distributions of normalized MED for the clusterings of sizes 2-25 found with the first iteration. The green line indicates the 12-clustering.

**Figure 9.**
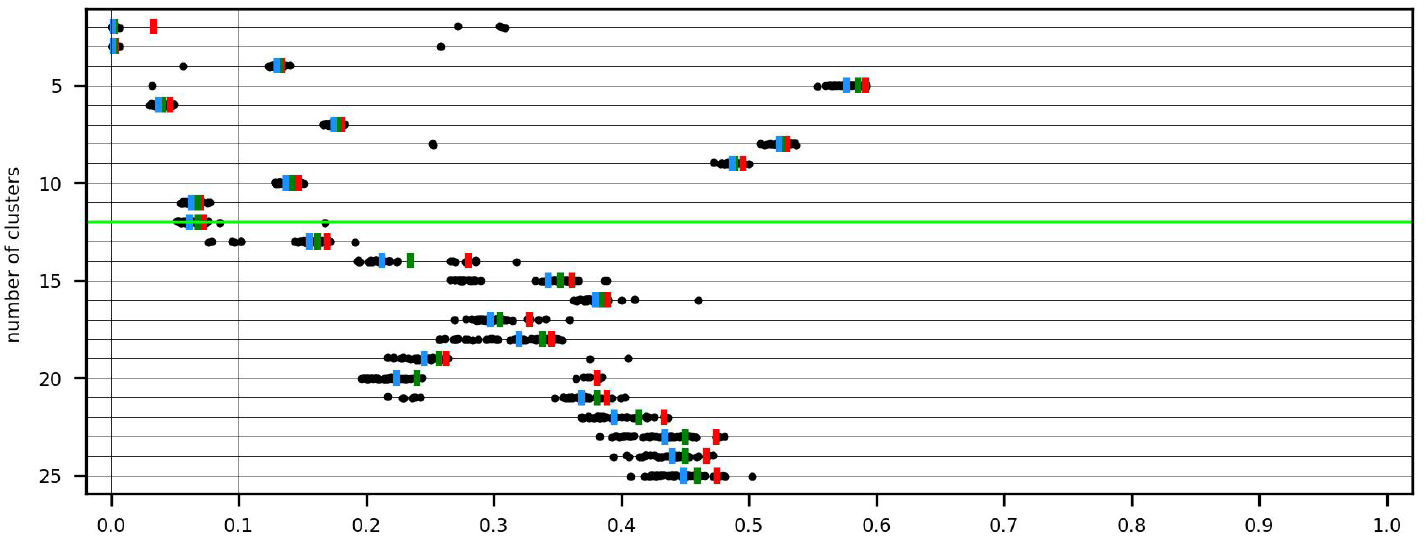
68k PBMC data (iteration 1): distributions of normalized MED for clusterings of sizes 2-25

Figure 10 displays the distributions of normalized CMER for the 12-clustering. The two smallest clusters, 8 and 11, are unstable. Their data lines are highlighted red because CMER=1 with all samples. The remaining clusters are stable. The clustering is admissible for downstream analysis because the unstable clusters have fewer than 500 cells. The blue, green, and red lines extending from the bottom to the top of the plot indicate the median, 75^*th*^, and 90^*th*^ percentiles of MED for the clustering.

**Figure 10.**
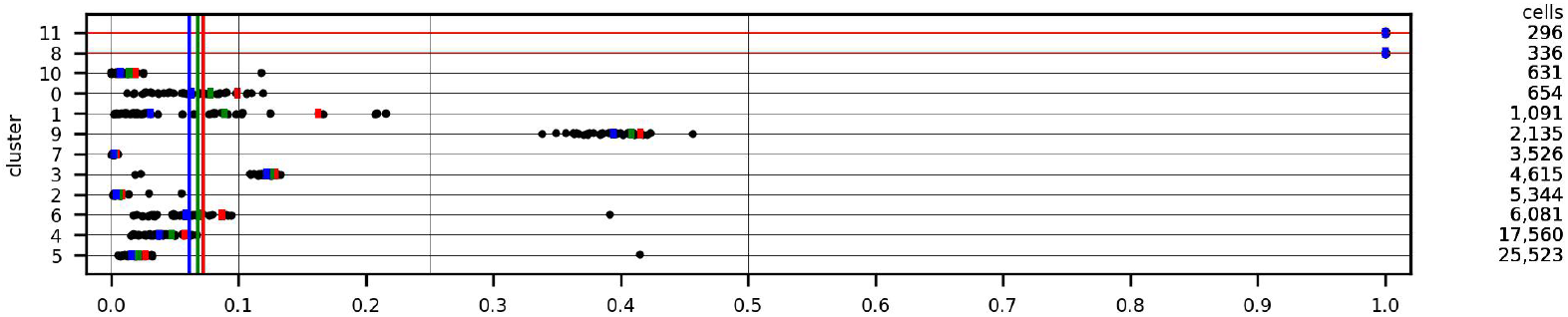
68k PBMC data (iteration 1): distributions of normalized CMER for the 12-clustering

Cell assignments to the ten k-means clusters summarized in Figure 3 of the paper [17] are evidently not publicly available. To attempt to compare clusterings found by our method with published results, the input data (68,579 cells) were clustered (k-means) using code from a 10x Genomics Github repository (see the section “Data availability” below). The sizes of the 10 clusters we obtained do not agree with the percentages in Figure 3b (nor is it obvious how these clusters correspond to the ones discussed in the paper). In particular, we obtained one small cluster – of 176 cells. The remaining 9 clusters range in size from three thousand to eighteen thousand cells. When the clusters obtained with our process were matched to these, none of the 176 cells in the smallest cluster were included – filtering discarded all of them.

Table 12 compares the 9 k-means clusters with the 12 hierarchical clusters. The adjusted Rand index equals 0.55.

- The majority of cells in k-means cluster 10 are split three ways. Most are in hierarchical cluster 7, which is stable. Approximately 300 cells each are in clusters 8 and 11, which are unstable.
- K-means cluster 9 is split four ways. Cells are divided among hierarchical clusters 0,1,9, and 10. Differential expression may help determine if these clusters are meaningful or spurious.
- K-means clusters 6 and 1 correspond to hierarchical clusters 2 and 3, respectively.
- 60% of the cells of k-means cluster 5 account for 80% of hierarchical cluster 6.
- 95% of cells in k-means cluster 4 are grouped with 90% of the cells in k-means cluster 7 in hierarchical cluster 5.
- 90% of cells in k-means cluster 2 are combined with nearly all of k-means cluster 3 and cells from other clusters into hierarchical cluster 4.

**Table 12.** 68k PBMC data (iteration 1): compare 9 k-means clusters with the 12-clustering.

| k-means cluster | hierarchical cluster |  |  |  |  |  |  |  |  |  |  |  | Total |
| --- | --- | --- | --- | --- | --- | --- | --- | --- | --- | --- | --- | --- | --- |
|  | 0 | 1 | 2 | 3 | 4 | 5 | 6 | 7 | 8 | 9 | 10 | 11 |  |
| 6 | 0 | 0 | 5,211 | 274 | 2 | 0 | 827 | 12 | 0 | 4 | 0 | 0 | 6,330 |
| 1 | 0 | 0 | 32 | 3,871 | 0 | 0 | 1 | 2 | 0 | 1 | 0 | 0 | 3,907 |
| 2 | 0 | 0 | 2 | 3 | 10,638 | 1,598 | 69 | 31 | 0 | 88 | 0 | 1 | 12,430 |
| 3 | 0 | 0 | 10 | 4 | 2,848 | 0 | 60 | 4 | 3 | 25 | 3 | 2 | 2,959 |
| 4 | 0 | 0 | 0 | 1 | 1,010 | 17,193 | 2 | 19 | 0 | 10 | 0 | 1 | 18,236 |
| 7 | 0 | 0 | 0 | 3 | 484 | 6,724 | 24 | 6 | 2 | 11 | 0 | 0 | 7,254 |
| 5 | 0 | 0 | 89 | 454 | 2,577 | 8 | 5,098 | 17 | 0 | 9 | 0 | 1 | 8,253 |
| 10 | 0 | 0 | 0 | 3 | 1 | 0 | 0 | 3,435 | 331 | 7 | 19 | 291 | 4,087 |
| 9 | 654 | 1,091 | 0 | 2 | 0 | 0 | 0 | 0 | 0 | 1,980 | 609 | 0 | 4,336 |
| Total | 654 | 1,091 | 5,344 | 4,615 | 17,560 | 25,523 | 6,081 | 3,526 | 336 | 2,135 | 631 | 296 | 67,792 |

We review a second analysis of the 68k PBMC data because it yields a clustering more compatible with the k-means clusters. It is the result of the third iteration described in section 2.8. After deleting outliers, 63,281 cells were retained. Figure 11 displays the distributions of normalized MED. Only the 2 and 3-clusterings are stable.

**Figure 11.**
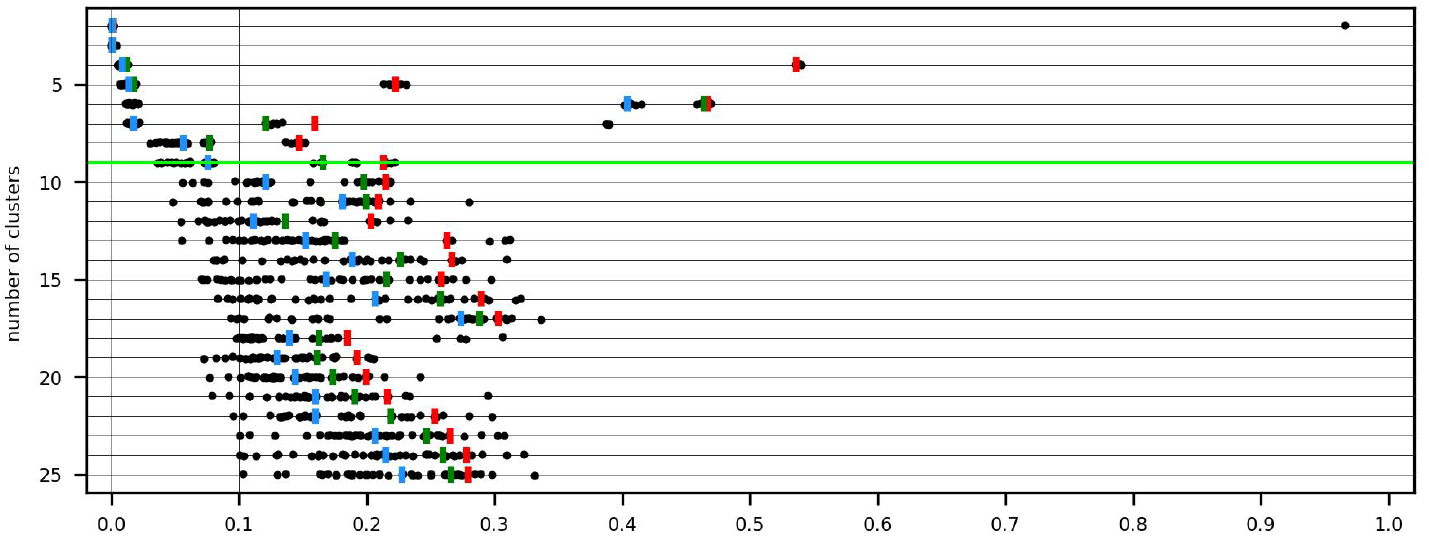
68k PBMC data (iteration 3): distributions of normalized MED for clusterings of sizes 2-25

Preliminary exploratory analysis using the *median* of MED instead of the 90^*th*^ percentile to define stable clusterings led to consideration of the 9-clustering (green highlighted line) because it is the largest with median MED (blue marker) less than or equal to 0.10. It fails to satisfy the criterion we now propose to define a stable clustering. The 90^*th*^ percentile of MED equals 0.21.

The confusion matrices in Tables 2 and 3 are for two of the samples summarized in Figure 12, which displays the distributions of normalized CMER for the 9 clusters.

**Figure 12.**
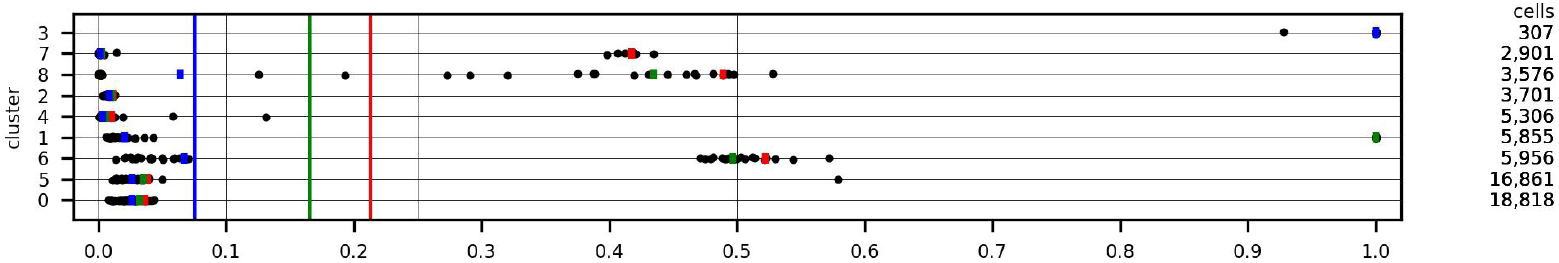
68k PBMC data (iteration 3): distributions of normalized CMER for the 9-clustering

Cluster 3 (top line) is unstable. CMER=1 with 39 samples. Cluster 1 (6^*th*^ line) is also unstable. The 75^*th*^ percentile of normalized CMER equals 1. Cluster 6 is unstable by a narrow margin. The remaining clusters are stable.

Table 13 compares the 9 k-means clusters with the 9 hierarchical clusters. The adjusted Rand index equals 0.66.

**Table 13.** 68k PBMC data (iteration 3): compare 9 k-means clusters with the 9-clustering.

| k-means cluster | hierarchical cluster |  |  |  |  |  |  |  |  | Total |
| --- | --- | --- | --- | --- | --- | --- | --- | --- | --- | --- |
|  | 0 | 1 | 2 | 3 | 4 | 5 | 6 | 7 | 8 |  |
| 4 | 16,371 | 298 | 1 | 0 | 0 | 1,042 | 3 | 10 | 19 | 17,744 |
| 7 | 1,058 | 5,499 | 1 | 0 | 0 | 481 | 25 | 13 | 5 | 7,082 |
| 1 | 0 | 0 | 3,086 | 286 | 28 | 0 | 1 | 0 | 0 | 3,401 |
| 6 | 0 | 0 | 173 | 20 | 5,167 | 2 | 591 | 0 | 5 | 5,958 |
| 2 | 1,388 | 51 | 3 | 0 | 0 | 10,477 | 65 | 86 | 23 | 12,093 |
| 3 | 0 | 0 | 5 | 0 | 7 | 2,725 | 49 | 15 | 3 | 2,804 |
| 5 | 1 | 7 | 432 | 1 | 104 | 2,134 | 5,222 | 5 | 14 | 7,920 |
| 9 | 0 | 0 | 0 | 0 | 0 | 0 | 0 | 2,772 | 0 | 2,772 |
| 10 | 0 | 0 | 0 | 0 | 0 | 0 | 0 | 0 | 3,507 | 3,507 |
| Total | 18,818 | 5,855 | 3,701 | 307 | 5,306 | 16,861 | 5,956 | 2,901 | 3,576 | 63,281 |

- Most of the cells in the unstable cluster 3 belong to k-means cluster 1. Recall that almost all of the cells in the two unstable clusters of the 12-clustering belong to k-means cluster 10.
- Each k-means cluster except 3 is unambiguously associated with a hierarchical cluster.

#### 25k retinal

The counts downloaded from the Gene Expression Omnibus contain data for 49,300 cells. We followed Lause et al. [18] by restricting to replicates p1 and r4-r6, netting counts for 22,292 genes and 24,769 cells. Thirty-nine cell clusters were reported (Figure 5D of [19]). Restricted to the cells we analyzed, cluster sizes range from 14 to 15,709 cells.

Filtering to exclude genes with nonzero counts on fewer than 50 cells retained 13,552 genes. Because data were not input through Seurat, cells were not screened for high mitochondrial DNA levels. 1,081 genes were found to be highly variable in the full data set and in all 40 samples.

The rank of the Pearson residuals matrix was estimated as 51 by optht. After mapping to a 51-dimensional SVD representation and excluding kNN outliers, 24,101 cells were retained. These cells belong to 37 of the 39 reported clusters.

Our objective was to evaluate clusterings of sizes 2-70 – the largest being greater than the number of reported cell clusters (39). The stopping conditions limited the largest clustering for at least one sample to a smaller size: 61 for iteration 1, 62 for iteration 2, 58 for iteration 3.

The largest stable clusterings found in the first and second iterations have 5 clusters. The largest stable clustering found in the third iteration has 11. The third iteration retained 22,416 cells, which belong to 36 of the 39 published cell clusters.

Figure 13 displays the distributions of normalized MED for the clusterings of sizes 2-58 found with the third iteration.

**Figure 13.**
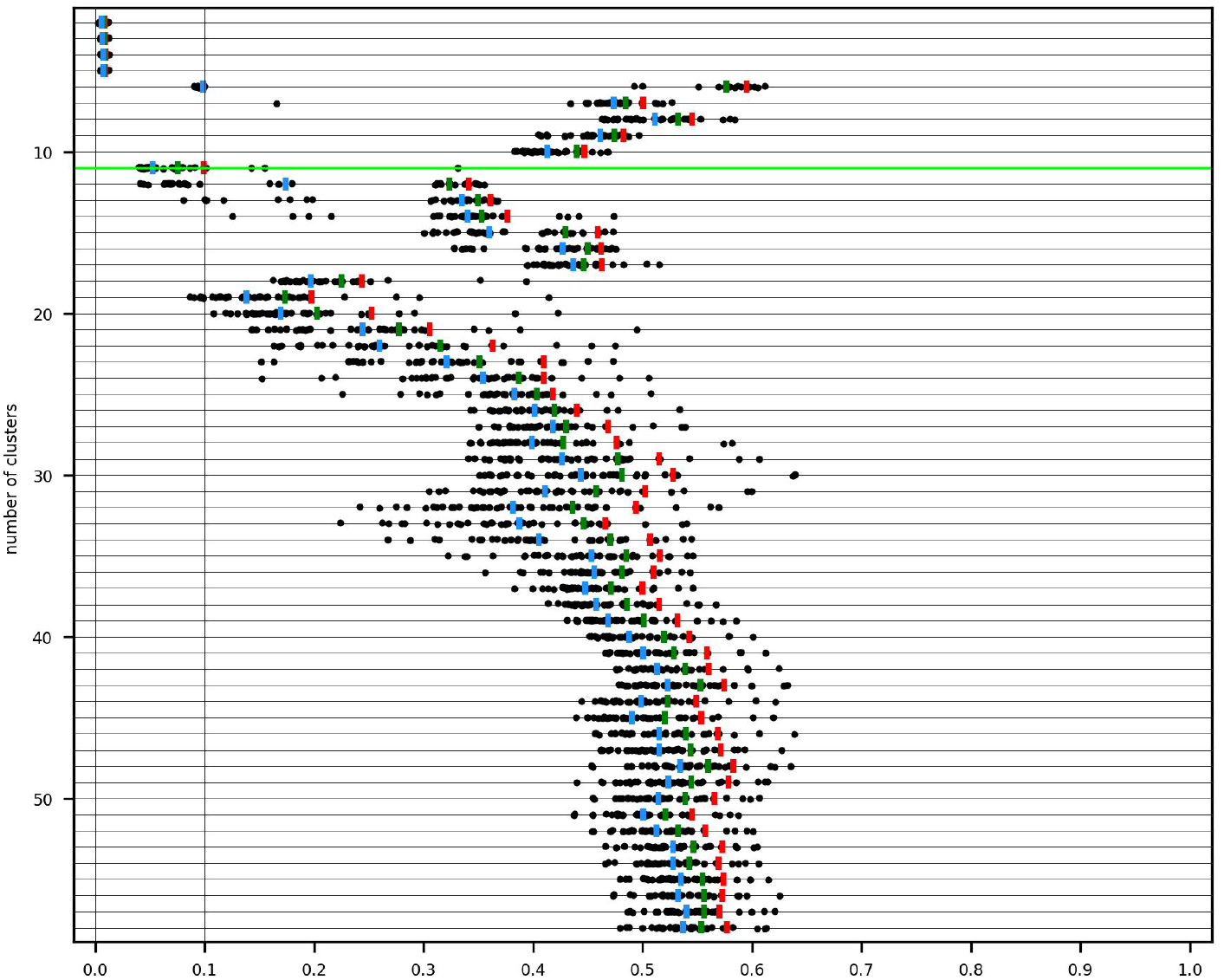
25k retinal data (iteration 3): distributions of normalized MED for clusterings of sizes 2-58

Figure 14 displays the distributions of normalized CMER for the 11 clusters. Cluster 9 is unstable. Because CMER=1 with all samples, the line is marked red. Cluster 6 is also unstable. The remaining clusters are stable. The clustering is admissible for downstream analysis.

**Figure 14.**
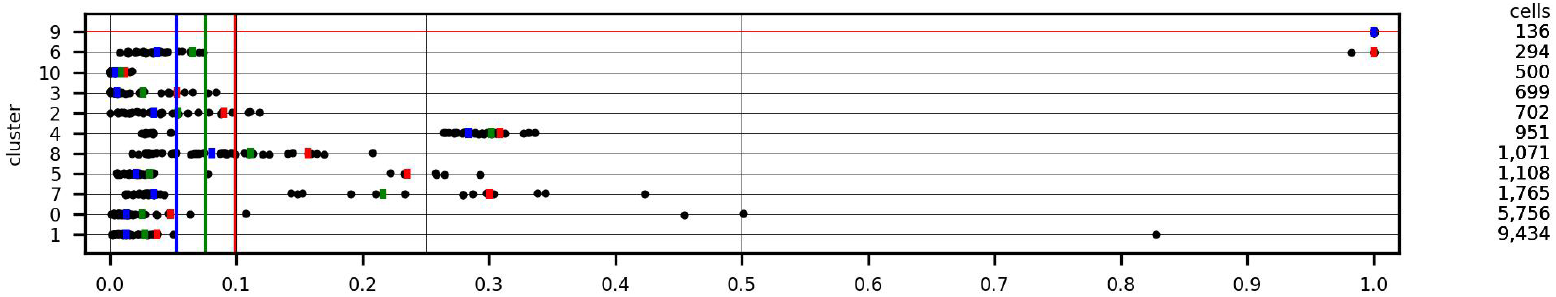
25k retinal data (iteration 3): distributions of normalized CMER for the 11-clustering

Table 14 illustrates the compatibility of the 11 hierarchical clusters with the reported cell clusters. The adjusted Rand index equals 0.49.

- Most members of cell cluster 24 (rods) belong to hierarchical clusters 0 and 1. We anticipate using differential expression analysis to evaluate this split.
- Similarly, cell cluster 26 is split between hierarchical clusters 2 and 3.
- Cell cluster 25 (cones) agrees closely with hierarchical cluster 8.
- Cell cluster 3 agrees almost exactly with the unstable hierarchical cluster 9.
- 80% of the cells in cluster 27 belong to the unstable hierarchical cluster 6.
- Cell cluster 34 agrees closely with the very stable hierarchical cluster 10.

**Table 14.**
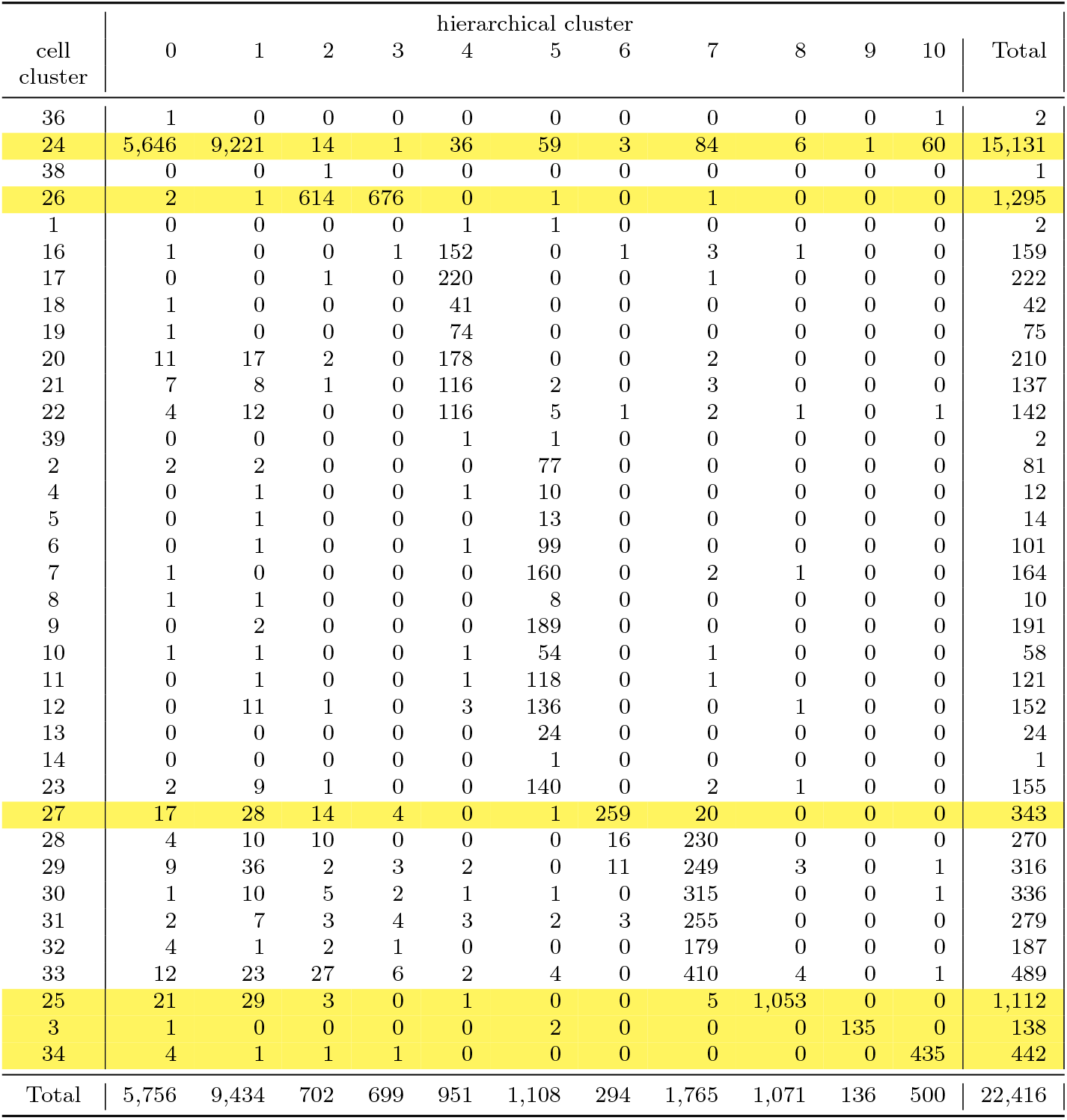
25k retinal data (iteration 3): compare 36 of 39 published clusters with the 11-clustering.

| cell<br>cluster | hierarchical cluster |  |  |  |  |  |  |  |  |  |  | Total |
| --- | --- | --- | --- | --- | --- | --- | --- | --- | --- | --- | --- | --- |
|  | 0 | 1 | 2 | 3 | 4 | 5 | 6 | 7 | 8 | 9 | 10 |  |
| 36 | 1 | 0 | 0 | 0 | 0 | 0 | 0 | 0 | 0 | 0 | 1 | 2 |
| 24 | 5,646 | 9,221 | 14 | 1 | 36 | 59 | 3 | 84 | 6 | 1 | 60 | 15,131 |
| 38 | 0 | 0 | 1 | 0 | 0 | 0 | 0 | 0 | 0 | 0 | 0 | 1 |
| 26 | 2 | 1 | 614 | 676 | 0 | 1 | 0 | 1 | 0 | 0 | 0 | 1,295 |
| 1 | 0 | 0 | 0 | 0 | 1 | 1 | 0 | 0 | 0 | 0 | 0 | 2 |
| 16 | 1 | 0 | 0 | 1 | 152 | 0 | 1 | 3 | 1 | 0 | 0 | 159 |
| 17 | 0 | 0 | 1 | 0 | 220 | 0 | 0 | 1 | 0 | 0 | 0 | 222 |
| 18 | 1 | 0 | 0 | 0 | 41 | 0 | 0 | 0 | 0 | 0 | 0 | 42 |
| 19 | 1 | 0 | 0 | 0 | 74 | 0 | 0 | 0 | 0 | 0 | 0 | 75 |
| 20 | 11 | 17 | 2 | 0 | 178 | 0 | 0 | 2 | 0 | 0 | 0 | 210 |
| 21 | 7 | 8 | 1 | 0 | 116 | 2 | 0 | 3 | 0 | 0 | 0 | 137 |
| 22 | 4 | 12 | 0 | 0 | 116 | 5 | 1 | 2 | 1 | 0 | 1 | 142 |
| 39 | 0 | 0 | 0 | 0 | 1 | 1 | 0 | 0 | 0 | 0 | 0 | 2 |
| 2 | 2 | 2 | 0 | 0 | 0 | 77 | 0 | 0 | 0 | 0 | 0 | 81 |
| 4 | 0 | 1 | 0 | 0 | 1 | 10 | 0 | 0 | 0 | 0 | 0 | 12 |
| 5 | 0 | 1 | 0 | 0 | 0 | 13 | 0 | 0 | 0 | 0 | 0 | 14 |
| 6 | 0 | 1 | 0 | 0 | 1 | 99 | 0 | 0 | 0 | 0 | 0 | 101 |
| 7 | 1 | 0 | 0 | 0 | 0 | 160 | 0 | 2 | 1 | 0 | 0 | 164 |
| 8 | 1 | 1 | 0 | 0 | 0 | 8 | 0 | 0 | 0 | 0 | 0 | 10 |
| 9 | 0 | 2 | 0 | 0 | 0 | 189 | 0 | 0 | 0 | 0 | 0 | 191 |
| 10 | 1 | 1 | 0 | 0 | 1 | 54 | 0 | 1 | 0 | 0 | 0 | 58 |
| 11 | 0 | 1 | 0 | 0 | 1 | 118 | 0 | 1 | 0 | 0 | 0 | 121 |
| 12 | 0 | 11 | 1 | 0 | 3 | 136 | 0 | 0 | 1 | 0 | 0 | 152 |
| 13 | 0 | 0 | 0 | 0 | 0 | 24 | 0 | 0 | 0 | 0 | 0 | 24 |
| 14 | 0 | 0 | 0 | 0 | 0 | 1 | 0 | 0 | 0 | 0 | 0 | 1 |
| 23 | 2 | 9 | 1 | 0 | 0 | 140 | 0 | 2 | 1 | 0 | 0 | 155 |
| 27 | 17 | 28 | 14 | 4 | 0 | 1 | 259 | 20 | 0 | 0 | 0 | 343 |
| 28 | 4 | 10 | 10 | 0 | 0 | 0 | 16 | 230 | 0 | 0 | 0 | 270 |
| 29 | 9 | 36 | 2 | 3 | 2 | 0 | 11 | 249 | 3 | 0 | 1 | 316 |
| 30 | 1 | 10 | 5 | 2 | 1 | 1 | 0 | 315 | 0 | 0 | 1 | 336 |
| 31 | 2 | 7 | 3 | 4 | 3 | 2 | 3 | 255 | 0 | 0 | 0 | 279 |
| 32 | 4 | 1 | 2 | 1 | 0 | 0 | 0 | 179 | 0 | 0 | 0 | 187 |
| 33 | 12 | 23 | 27 | 6 | 2 | 4 | 0 | 410 | 4 | 0 | 1 | 489 |
| 25 | 21 | 29 | 3 | 0 | 1 | 0 | 0 | 5 | 1,053 | 0 | 0 | 1,112 |
| 3 | 1 | 0 | 0 | 0 | 0 | 2 | 0 | 0 | 0 | 135 | 0 | 138 |
| 34 | 4 | 1 | 1 | 1 | 0 | 0 | 0 | 0 | 0 | 0 | 435 | 442 |
| Total | 5,756 | 9,434 | 702 | 699 | 951 | 1,108 | 294 | 1,765 | 1,071 | 136 | 500 | 22,416 |

#### 65k lung

The paper [21] describes separately clustering the data for each patient. Our analysis included batch correction. Each patient’s data were treated as an independent batch.

The data set contains counts for 26,485 genes and 65,662 cells. The accompanying metadata file provides 57 cell type annotations.

Excluding data for blood cells retained 60,993 cells. Filtering to exclude genes with nonzero counts on fewer than 50 cells retained 17,470 genes. Because data were not input through Seurat, cells were not screened for high mitochondrial DNA levels. 1,659 genes were found to be highly variable in the full set of data and all 40 samples.

The rank of the Pearson residuals matrix was estimated as 305 by optht. After mapping to a 305-dimensional SVD representation and excluding kNN outliers, 60,114 cells were retained.

Our objective was to evaluate clusterings of sizes 2-70 – the largest being greater than the number of reported cell types (57). The stopping conditions limited the largest clustering for at least one sample to a smaller size: 69 for iteration 1, 48 for iteration 2, 41 for iteration 3. The largest stable clustering found in the first iteration has 19 clusters. The largest stable clusterings found in the second and third iterations have 13 clusters.

Figure 15 displays the distributions of normalized MED for the clusterings of sizes 2-69 found with the first iteration. The green lines highlight data for the clusterings of sizes 16 and 19, which are reviewed.

**Figure 15.**
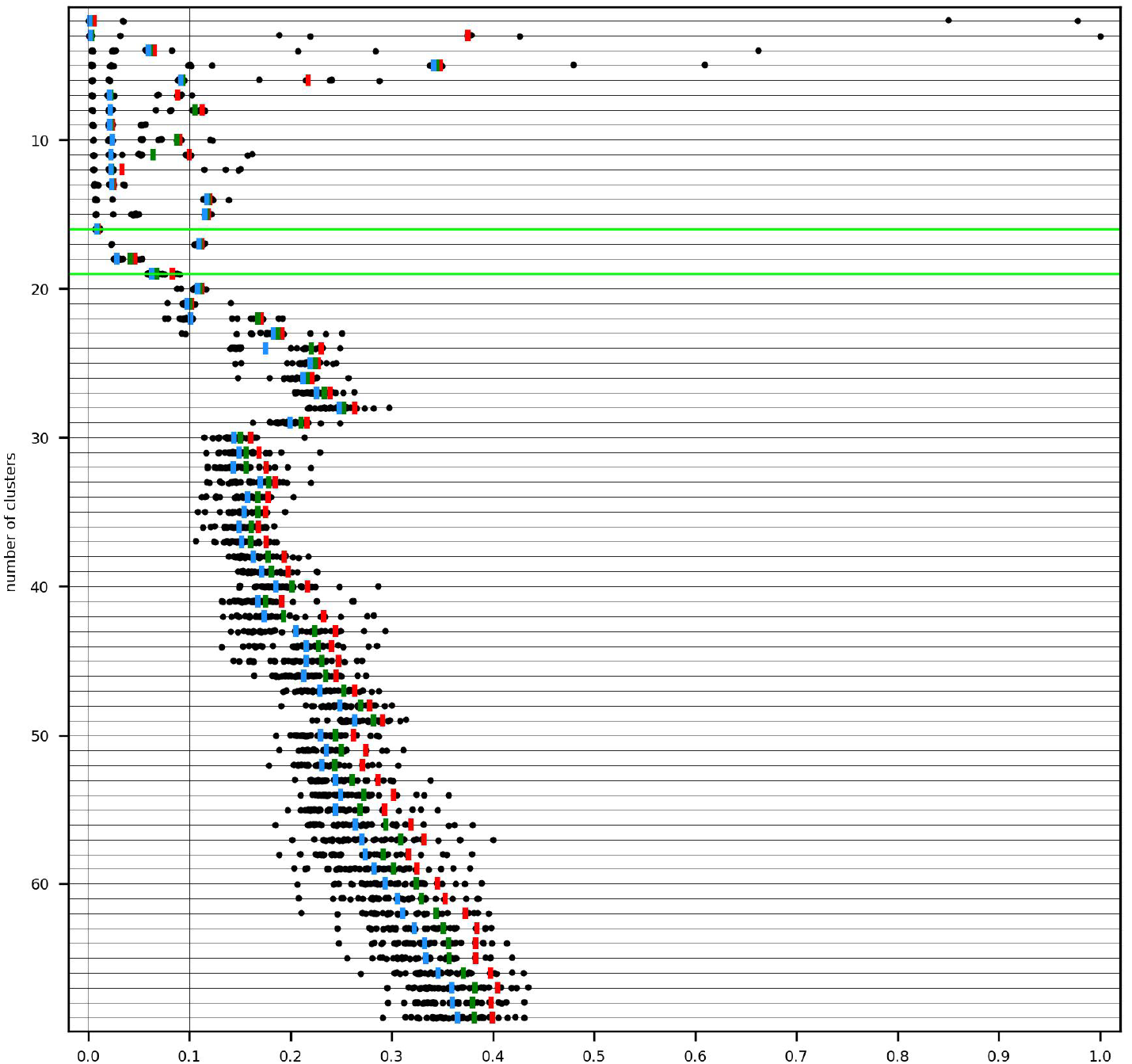
65k lung data (iteration 1): distributions of normalized MED for clusterings of sizes 2-69

Figure 16 displays the distributions of normalized CMER for the 19 clusters. Clusters 1 and 4 (first and third lines) are unstable. The remaining clusters are stable. The clustering is admissible for downstream analysis.

**Figure 16.**
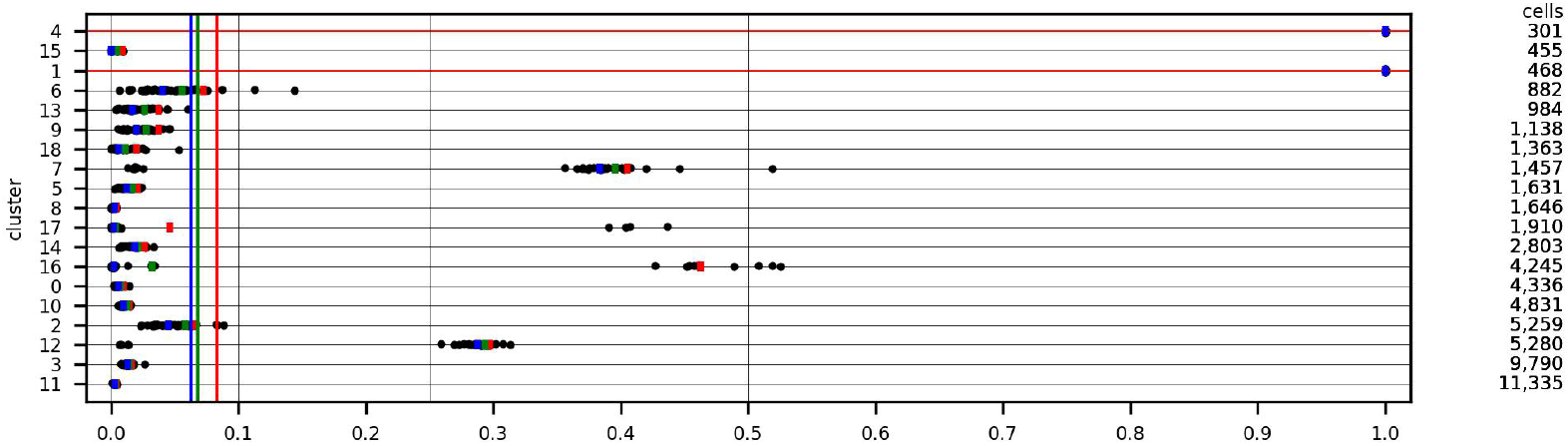
65k lung data (iteration 1): distributions of normalized CMER for the 19-clustering

Because blood cells were excluded from our analysis, data for one cell type were eliminated. Table 15 illustrates the compatibility of the 19 hierarchical clusters with the 56 retained cell types. The adjusted Rand index equals 0.65. The majority of macrophages are divided between clusters 2 and 3.

**Table 15.**
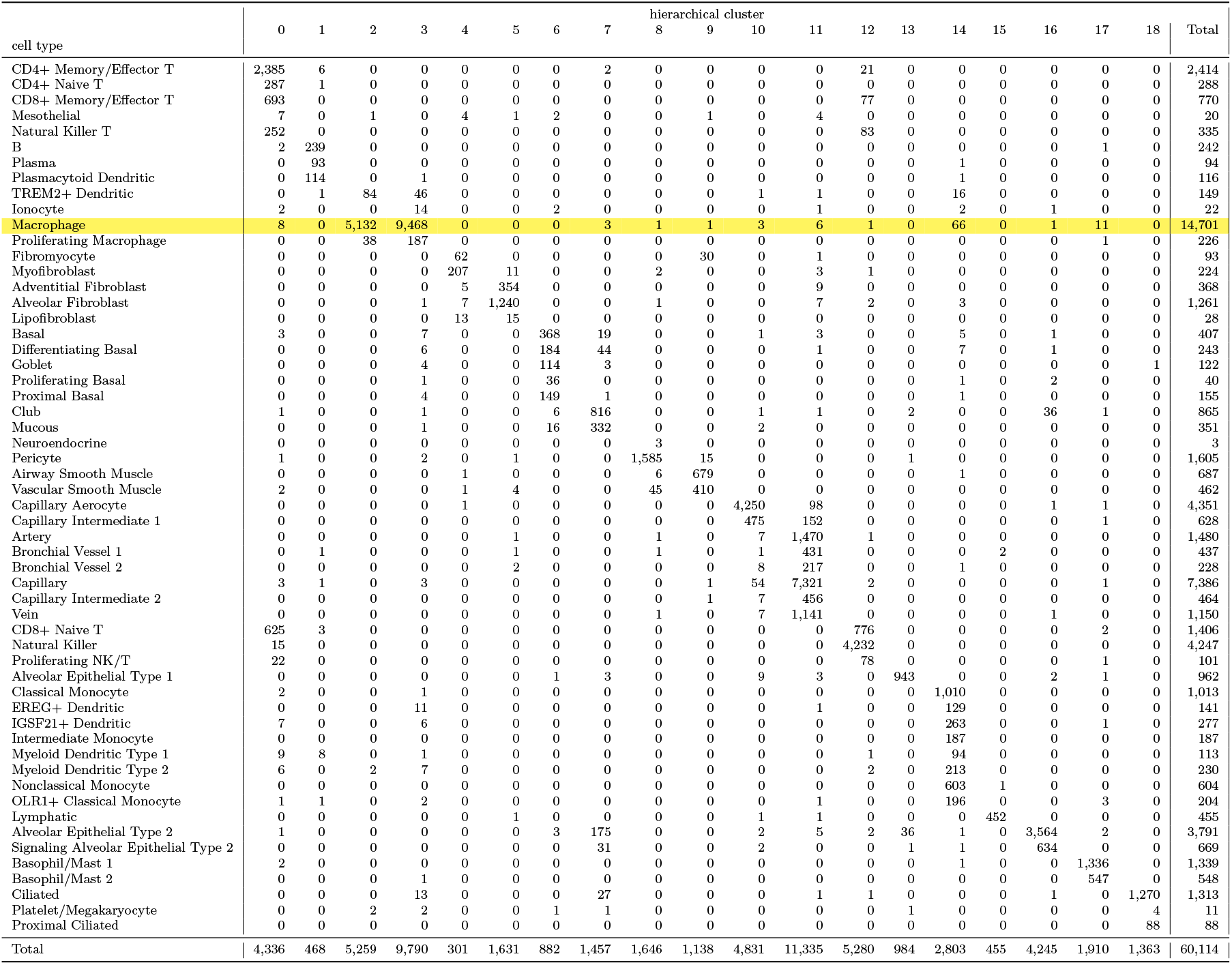
65k lung data (iteration 1): compare 56 of 57 reported cell types with the 19-clustering.

The 16-clustering of the same data is reviewed because the results – for both the clustering and its clusters – are the most stable found in the four large data sets. Referring to Figure 15, the *maximum* value of normalized MED equals 0.01. We cannot explain this why it is exceptionally small.

Figure 17 displays the distributions of normalized CMER for the 16 clusters. All are stable.

**Figure 17.**
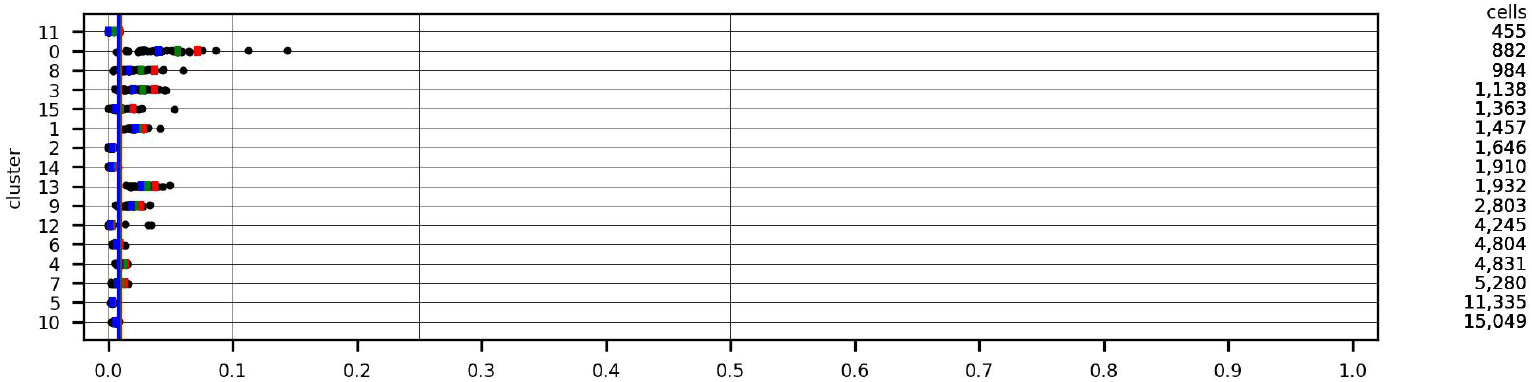
65k lung data (iteration 1): distributions of normalized CMER for the 16-clustering

Table 16 compares the published cell types with the 16 hierarchical clusters. The adjusted Rand index equals 0.81. The majority of macrophages belong to cluster 10.

**Table 16.**
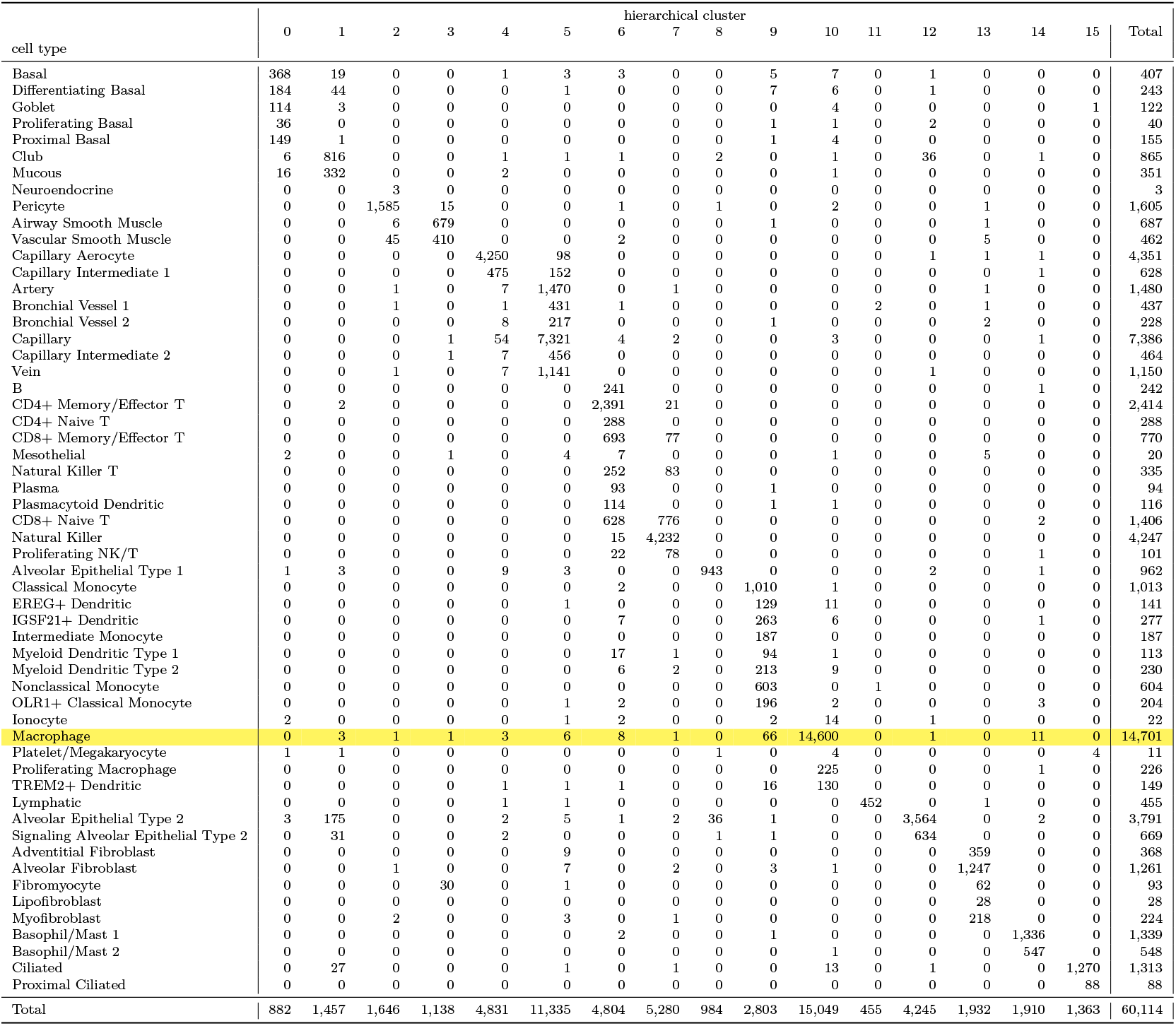
65k lung data (iteration 1): compare 56 of 57 reported cell types with the 16-clustering.

#### 100k breast cancer

Discussion in [23] and programs posted at [38] suggest that batch correction was performed for each patient’s data for some, if not all, analyses. In our analysis, each patient’s data were treated as an independent batch.

The data set contains counts for 29,733 genes and 100,064 cells. Nine major cell types were reported, as well as 29 minor cell types and 49 cell type subsets.

Filtering to exclude genes with nonzero counts on fewer than 50 cells retained 21,354 genes. The data include cells for which up to 20% of counts are due to mitochondrial DNA. (The percentage exceeds 5% for half of the cells.) 1,704 genes were found to be highly variable in the set of counts for all cells and in all 40 samples.

The rank of the Pearson residuals matrix was estimated as 434 by optht. After mapping to a 434-dimensional SVD representation and excluding kNN outliers, 98,681 cells were retained.

Our objective was to evaluate clusterings of sizes 2-70 – the largest being greater than the number of reported cell type subsets (49). The stopping conditions limited the largest clustering for at least one sample to a smaller size: 58 for iteration 1, 39 for iteration 2, 51 for iteration 3. No stable clusterings were found. The smallest values found for the 90^*th*^ percentile of MED are 0.15 (first iteration), 0.16 (second), and 0.107 (third).

Figure 18 displays the distributions of normalized MED for the clusterings of sizes 2-51 found with the third iteration. We review the 9-clustering, which has the smallest 90^*th*^ percentile (0.107). This near miss from being judged stable illustrates a disadvantage of hard thresholds. We describe the clustering as *marginally unstable*.

**Figure 18.**
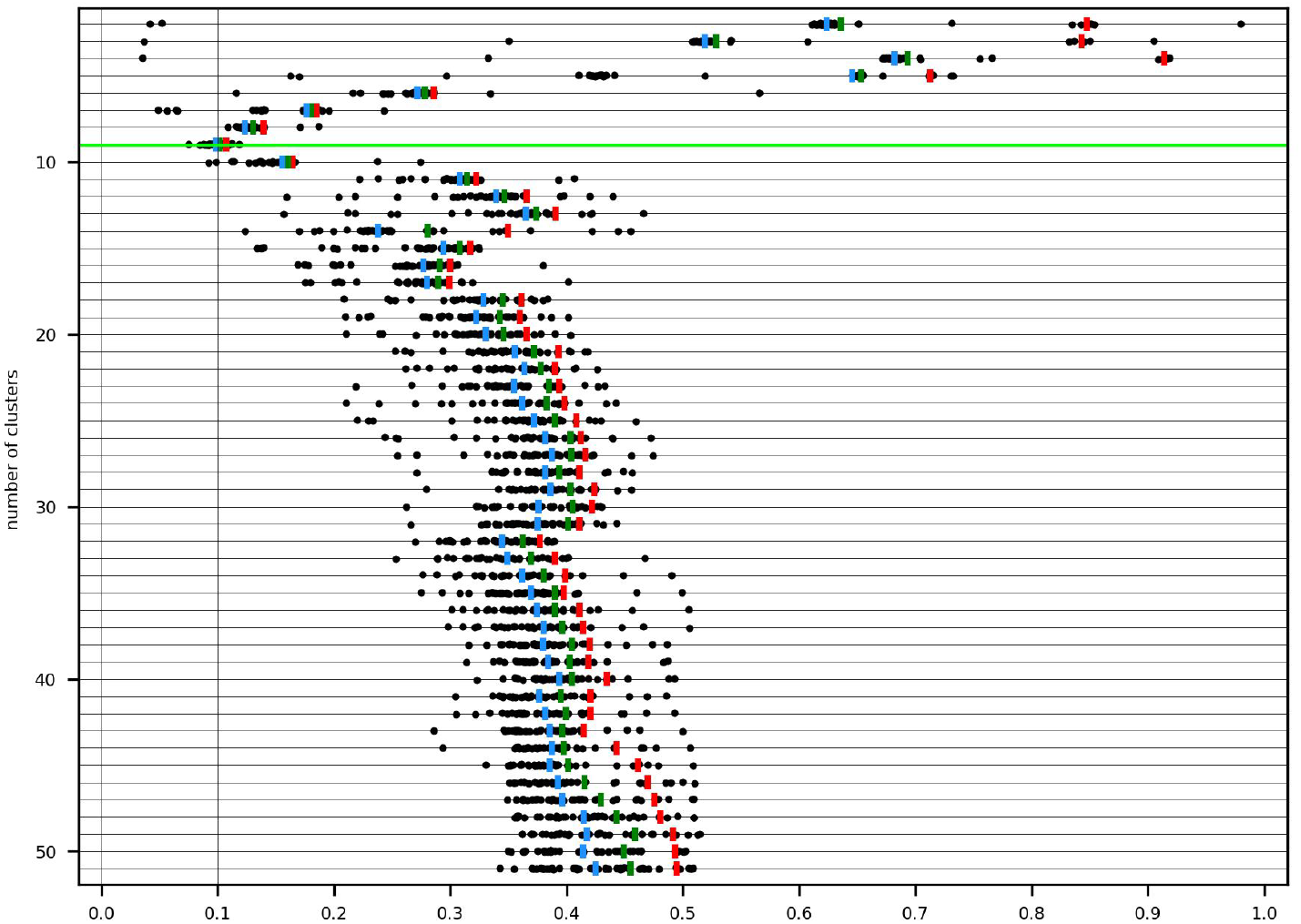
100k breast cancer data (iteration 3): distributions of normalized MED for clusterings of sizes 2-51

Figure 19 displays the distributions of normalized CMER for the 9 clusters. Cluster 6 (top line) is unstable. CMER equals 1 with 39 of the 40 samples. Cluster 0 (fifth line) is also unstable. The 90^*th*^ percentile of CMER equals 0.55, larger than the threshold of 0.50 that defines stable clusters. This is a second reminder of the disadvantage of hard thresholding. The remaining clusters are stable. The clustering is not admissible for downstream analysis.

**Figure 19.**
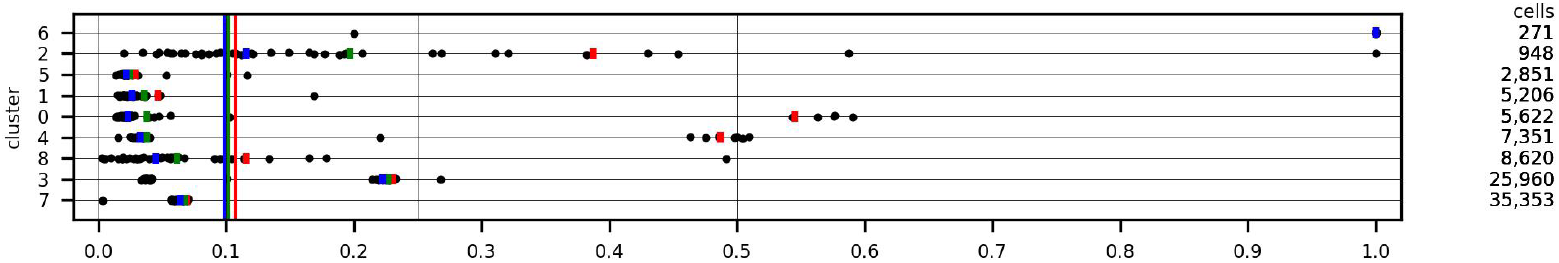
100k breast cancer data (iteration 3): distributions of normalized CMER for the 9-clustering.

The third iteration retained 92,182 cells, which include all 49 published cell type subsets. Filtering had a disproportionate impact on plasmablasts, one of the 9 major cell types. The data downloaded from the GEO included counts for 100,064 cells (Table 1) of which 3,524 are plasmablasts. Excluding kNN outliers before clustering retained 98,681 cells (Table 4) of which 2,290 are plasmablasts. They account for 1,234 of the 1,383 cells excluded as outliers in the first iteration. Only 801 plasmablasts were retained in the third iteration.

Table 17 compares the 9 hierarchical clusters with the major cell types. The adjusted Rand index equals 0.86.

- The hierarchical clusters do not separate normal from cancer epithelial cells, though approximately a quarter of the normal cells are assigned to cluster 2, accounting for more than 95% of its membership.
- The unstable cluster 6 represents a segment of myeloid cells.
- 90% of the plasmablasts that survived the filtering iterations are split between clusters 3 and 5.

**Table 17.** 100k breast cancer data (iteration 3): compare major cell types with the 9-clustering.

| major cell type | hierarchical cluster |  |  |  |  |  |  |  |  | Total |
| --- | --- | --- | --- | --- | --- | --- | --- | --- | --- | --- |
|  | 0 | 1 | 2 | 3 | 4 | 5 | 6 | 7 | 8 |  |
| CAFs | 5,364 | 157 | 2 | 73 | 4 | 1 | 0 | 8 | 13 | 5,622 |
| PVL | 39 | 4,915 | 4 | 46 | 26 | 0 | 0 | 2 | 4 | 5,036 |
| Cancer Epithelial | 165 | 61 | 19 | 21,868 | 18 | 23 | 0 | 1,402 | 144 | 23,700 |
| Normal Epithelial | 1 | 3 | 909 | 2,910 | 1 | 2 | 0 | 22 | 10 | 3,858 |
| Plasmablasts | 6 | 9 | 3 | 431 | 43 | 300 | 2 | 7 | 0 | 801 |
| Endothelial | 12 | 22 | 0 | 59 | 7,109 | 2 | 0 | 3 | 4 | 7,211 |
| B-cells | 3 | 14 | 8 | 254 | 98 | 2,425 | 3 | 25 | 45 | 2,875 |
| T-cells | 4 | 7 | 2 | 205 | 12 | 85 | 0 | 33,831 | 13 | 34,159 |
| Myeloid | 28 | 18 | 1 | 114 | 40 | 13 | 266 | 53 | 8,387 | 8,920 |
| Total | 5,622 | 5,206 | 948 | 25,960 | 7,351 | 2,851 | 271 | 35,353 | 8,620 | 92,182 |

Table 18 illustrates the compatibility of the 9 hierarchical clusters with the 49 cell type subsets identified in the paper. The adjusted Rand index equals 0.20.

- Cluster 2, mostly normal epithelial cells, corresponds to myoepithelial cells.
- Cluster 6 (unstable) corresponds to myeloid c4 DCs pDC IRF7 cells.

**Table 18.**
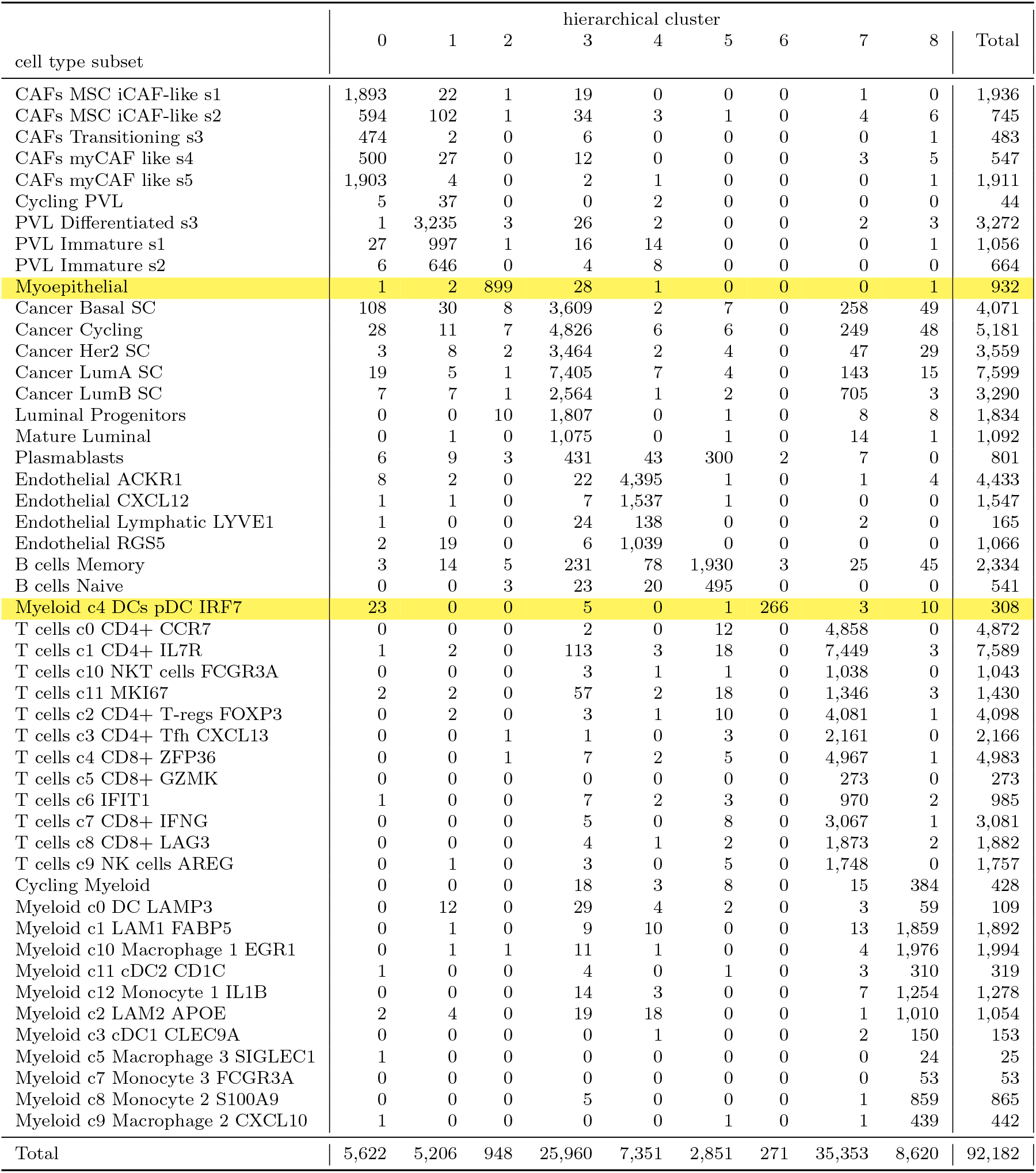
100k breast cancer data (iteration 3): compare cell type subsets with the 9-clustering.

## 4 Discussion and Conclusions

We have shown that it is analytically possible and computationally feasible to identify stringently-defined stable clusterings and clusters of scRNA-seq data presented as UMI counts for as many as 100 thousand cells.

Results for the retinal, lung, and breast cancer data suggest one direction for future work: using differential expression analysis to validate results. The published results identified large numbers of cell groups (39 for retinal, 57 for lung, 49 for breast cancer). Our method gave fewer (11, 19, and 9, respectively) though their compatibility (Tables 14, 15, 16, and 18) suggests a question. *If less stringent criteria (higher thresholds for MED and CMER) define a larger number of stable clusters, how can they be evaluated?* It may be sufficient to require that the differential expression of analysis genes – between pairs of stable clusters – be consistent across all samples. If a gene’s Pearson residuals are low on cluster *c*_1_ and high on *c*_2_ in the full set of cells, require that this hold with every sample, and that the estimates of the differences be consistent. Preliminary exploratory analysis suggests that this may not be satisfied by clusters with large CMER. Recall the third iterative analysis of the PBMC data. The 9-clustering was reviewed because of its greater compatibility with the k-means clustering produced by the 10x Genomics code (Table 13). CMER values of cluster 1 are large. Their 75^*th*^ percentile equals 1 (Figure 12). Comparing the distributions of Pearson residuals of analysis genes for cells in cluster 1 with cells in other clusters did not show the same consistency across samples that we found for clusters defined as stable with the criterion of section 2.1 – that the 90^*th*^ percentile of normalized CMER be less than or equal to 0.50.

This work complements that of Levine and Domany [1], who proposed evaluating a clustering’s stability by comparing it with clusterings of random samples.

These algorithmic features may be novel in the scRNA-seq context:

- quantifying stability by comparing a set of cells with pairs of random complementary (disjoint) samples, each containing approximately half of the cells
- using normalized MED to compare clusterings of samples with clusterings of the full set of cells
- using divisive hierarchical spectral clustering; defining the affinity of two points as the inverse of the distance between them may be novel in any context
having constructed a dendrogram, defining the distance from a node to its daughters as the normalized cut (the quantity minimized by spectral clustering), then using the distances from the root to the nodes to map the dendrogram to a set of nested clusterings

Returning to the relation between our clustering results and published work: the four clusters of the Zhengmix4eq data set agree very closely with the ground truth labels.

For the Zhengmix8eq data, the failure of our method to match all of the ground truth labels, specifically for three T-cell subtypes, seems to be typical. Lun [7] mentions “various flavours of T cells” as examples of “cell types that are difficult to separate.” The recent paper by Mullan et al. [39] is concerned with the failure of (unsupervised) clustering to correctly identify T-cell subtypes.

Results for the monocytes suggest that our approach will not find spurious clusters in homogeneous data.

The 11-clustering of the retinal data splits two of the published cell clusters: 24 (rods) and 26. Similarly, in the 19-clustering of the lung data, the majority of macrophages are divided between two hierarchical clusters. Future work will explore using differential expression to determine if these splits are meaningful.

Explaining the exceptional stability of the 16-clustering of the lung data may be essential to obtaining comparable results for other data.

Results for the breast cancer data prompt two observations. (1) In section 3.5 we noted that iterating to remove outliers has a disproportionate impact on plasmablasts. Understanding this might point to process improvements. (2) The 9-clustering is described as marginally unstable because the 90^*th*^ percentile of MED equals 0.107 – slightly larger than the threshold of 0.10. Similarly, cluster 0 is unstable because the 90^*th*^ percentile CMER equals 0.55 – greater than the threshold of 0.50. As an alternative to specifying thresholds for summary statistics of MED and CMER to define stability, their values could be reported without any judgment – like p-values for a statistical test. If future work shows that differential expression can validate the identification of stable clusterings and clusters, then MED and CMER thresholds may become less important. Statistics characterizing consistent differential expression might be alternative decision criteria.

Other issues may deserve attention:

- Defining a principled rule to decide when unstable clusters render a stable clustering inadmissible for downstream analysis. Requiring that unstable clusters have fewer than 500 cells is a simple expedient.
- More efficient calculation, or estimation, of the distances between points in the SVD representation. We used the scikit-learn euclidean distances function. It is slow for the lung and breast cancer data, with many points in high dimensions.
- Better understanding the iterative process to remove count outliers, because of its inconsistent impact. The third iteration gave the largest admissible clustering of the retinal data and the marginally unstable clustering of the breast cancer data. For the PBMC and lung data, the first iteration gave the largest admissible clusterings. It is not obvious how many iterations are appropriate, or if a point of diminishing returns can be found – to stop the process because subsequent iterations would not improve results. Three iterations were used for illustration. More were not performed because of the cost of calculating the distance between cells.
- Exploring whether – as speculated in section 2.8 – for a UMI count matrix to admit a stable clustering, it is necessary that the distributions of UMI counts (or of lower-dimension derived data such as the SVD representation) in all sufficiently large samples agree.

## Data availability

### Zhengmix4eq/8eq

Loaded from the Bioconductor DuoClustering2018 package [15]

### 68k PBMC

Downloaded from the 10x Genomics website https://www.10xgenomics.com/datasets/fresh-68-k-pbm-cs-donor-a-1-standard-1-1-0

Select ‘Explore Data’ then ‘Output and supplemental files.’ Specifying the gene/cell matrix (filtered) 124 MB file yields the fresh_68k_pbmc_donor_a_filtered_gene_bc_matrices.tar file, which contains the files matrix.mtx, barcodes.tsv, and genes.tsv with data for 32,738 genes and 68,579 cells

K-means clustering of the 68,579 cells was performed using an excerpt from the program main_process_68k_pbmc.R posted in https://github.com/10XGenomics/single-cell-3prime-paper/tree/master/pbmc68k_analysis

This program, along with the util.R utility program and two RDS files accessible with links in the README file have been placed in our Github repository (see Code availability below) in the sub-folder 10x_Genomics_material of the 68k_PBMC folder

### CD14 Monocytes

Downloaded from the 10x Genomics website

### 25k retinal

These instructions are from https://github.com/berenslab/umi-normalization, associated with [18]

Counts

- Visit https://www.ncbi.nlm.nih.gov/geo/
- Search for GSE63472
- Download GSE63472_P14Retina_merged_digital_expression.txt.gz (50.7 MB)
- Extract to GSE63472_P14Retina_merged_digital_expression.txt

Cluster annotations

- Download from http://mccarrolllab.org/wp-content/uploads/2015/05/retina_clusteridentities.txt

### 65k lung

Metadata krasnow_hlca_10x_metadata.csv and counts krasnow_hlca_10x_UMIs.csv were downloaded from https://www.synapse.org/#!Synapse:syn21041850. The count file is dense: it explicitly contains all zero counts

### 100k breast cancer

As documented in [23], data are available through the GEO Series accession number GSE176078. GSE176078_Wu_etal_2021_BRCA_scRNASeq.tar.gz contains the files count_matrix_sparse.mtx, count_matrix_barcodes.tsv, count_matrix_genes.tsv, and metadata.csv

## Code availability

The Github repository https://github.com/victorkleb/scRNA-seq stable clust contains 7 folders, one for each data set studied. Each folder contains the pipeline for analyses described in this article.

## Acknowledgments

The author thanks Martin Surks and Will Townes for helpful conversations and suggestions.

## Appendix A Closed-form expression for the mean sum of squares of Pearson residuals with a Poisson model

For convenience, this is copied – with minor changes – from [28]. The formula is sparse – involving only entries of the UMI count matrix and row and column sums. Hence, the SSQ of genes’ Pearson residuals can be computed sparsely; the dense Pearson residuals matrix is not needed.

We use the notations

*X*_*gc*_ = the UMI count for gene *g* in cell *c*

*G*_*g*_ = ∑_*c*_ *X*_*gc*_ the total count for gene *g*

*π*_*c*|*g*_ = *X*_*gc*_*/G*_*g*_ the fraction of the total count of gene *g* that is in cell *c*

*D*_*c*_ = ∑_*g*_*X*_*gc*_ the total count for cell *c* – its sequencing depth

*T* =∑_*gc*_ *X*_*gc*_ the total of all counts in the matrix

*C* = the total number of cells in the matrix

*D* = *T/C* the mean sequencing depth of the matrix

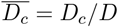 the normalized sequencing depth of cell *c*

The Pearson residuals are

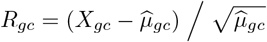

with the maximum likelihood estimate

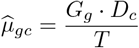

Squaring gives

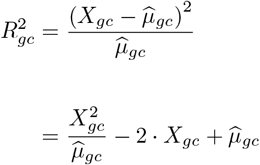

Summing over all cells *c* gives ***S***_*g*_, the sum of squares of Pearson residuals for gene *g*:

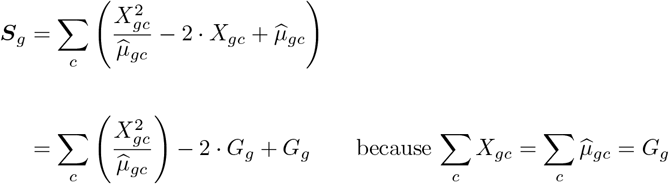

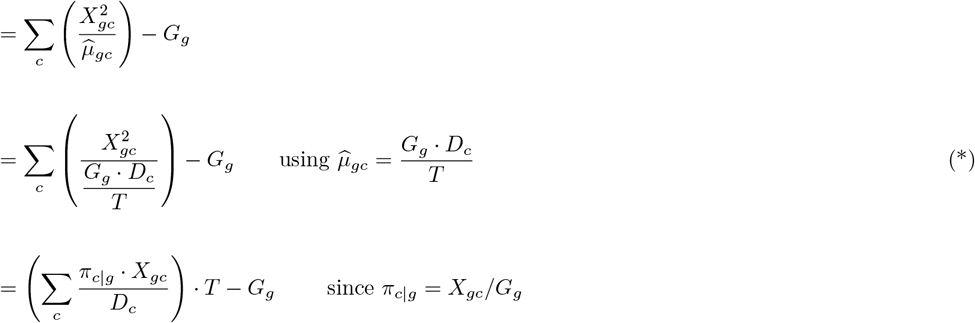

Dividing by the number of cells in the UMI matrix, *C*, gives the mean sum of squares of Pearson residuals for gene *g*:

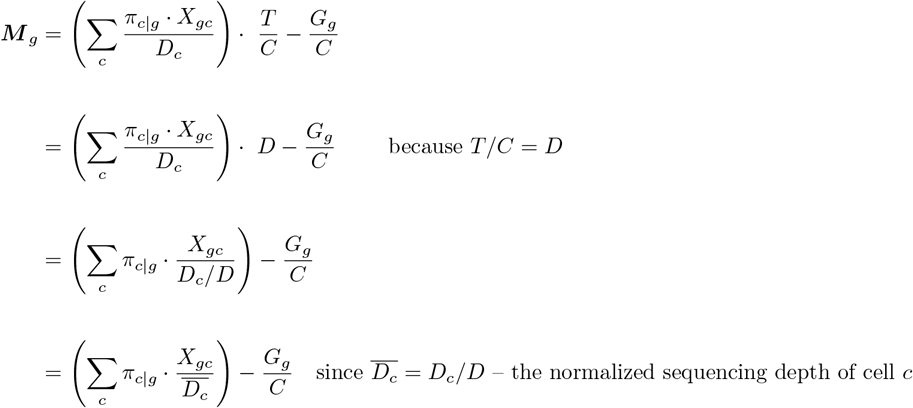

Observations:

- The summation includes *only* cells on which gene *g* has nonzero counts
- The mean value of the denominators 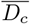 *over all cells in the UMI count matrix* is 1
- The subtracted term 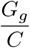 is negligible for lowly expressed genes
- Since ∑*π*_*c*|*g*_ = 1, the summation is the convex combination of the ratios 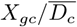. This provides intuition for the fact that many values of ***M*** _*g*_ are near 1. They are convex combinations of terms that are often close to 1: most nonzero counts *X*_*gc*_ equal 1 and the mean of the 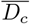 equals 1

## Appendix B Closed-form expression for a cell’s contribution to the sum of squares of a gene’s Pearson residuals

Continuing the above derivation from equation (*) identifies each cell’s contribution to *S*_*g*_

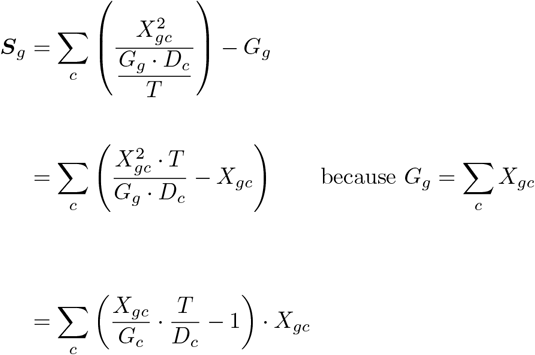

Denoting the contribution of cell *c* to ***S***_*g*_ as *K*_*gc*_

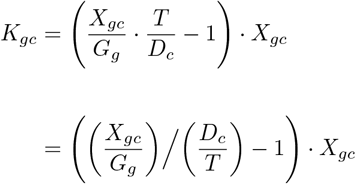

Observations:

- The numerator *X*_*gc*_*/G*_*g*_ is the fraction of the count of gene *g* in cell *c*
- The denominator *D*_*c*_*/T* is the fraction of the total UMI count in cell *c*
- Hence, the ratio compares the contribution of cell *c* to the count of *g* with the cell’s contribution to the matrix’s total count
- If this ratio equals 1, there is *no* contribution to the SSQ of the PR of *g*
- If it is less than 1, the cell’s contribution to the SSQ of the PR is *negative*

