## Supplement 2 for "Finding stable clusterings of single-cell RNA-seq data" (from another version of this preprint: version 2, 2026-01-18)

**Supplement 2: Distributions of normalized MED for clusterings of retinal, lung, and breast cancer data**

The distributions of MED displayed in the article are restricted to clusterings of sizes 2-25. No larger clusterings produced by our approach (restricting to clusterings of sizes 2-70) are stable. Because the figures including all generated clusterings are bulky, they are provided separately in this Supplement.

**25k retinal**

Although our objective was to evaluate clusterings of sizes 2-70, stopping conditions limited the largest clusterings found for all samples to size 61 for iteration 1; 62 for iteration 2; and 58 for iteration 3. Figures S2-1, S2-2, and S2-3 display the distributions of normalized MED for the three iterations.

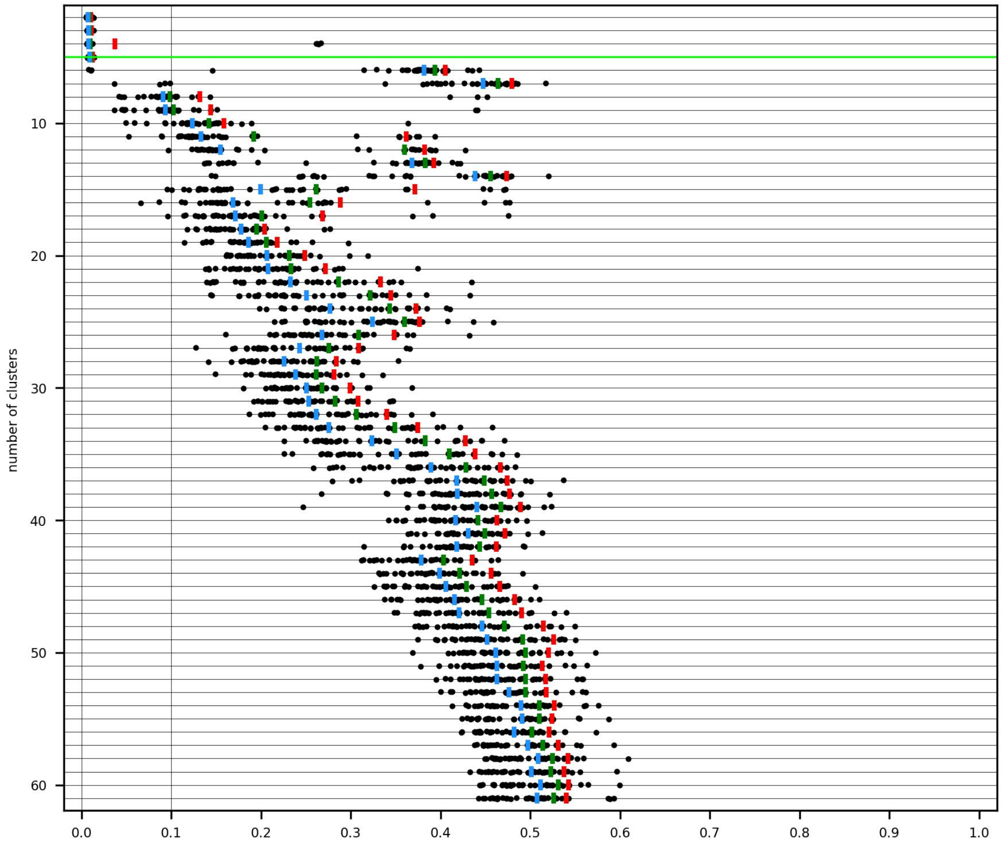


**Figure S2-1** 25k retinal data (iteration 1): distributions of normalized MED for all generated clusterings; the largest stable clustering has size 5 (green highlighted line)

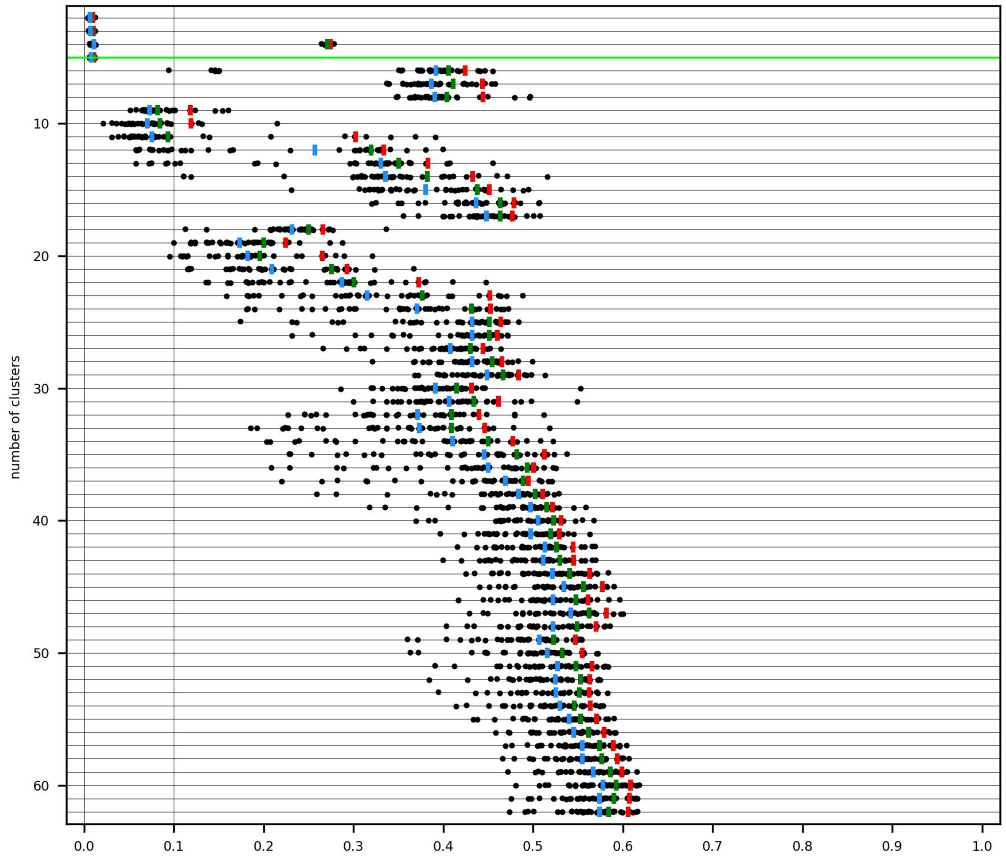


**Figure S2-2** 25k retinal data (iteration 2): distributions of normalized MED for all generated clusterings; the largest stable clustering has size 5

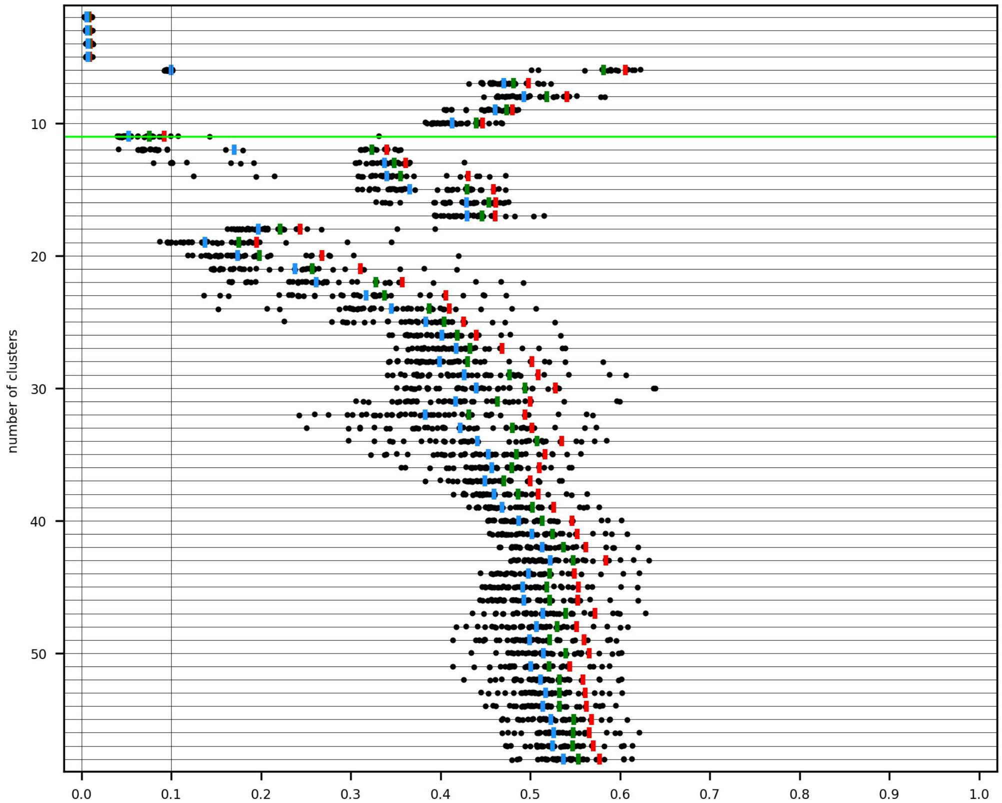


**Figure S2-3** 25k retinal data (iteration 3): distributions of normalized MED for all generated clusterings; the largest stable clustering has size 11

### 65k lung

As noted for the retinal data, our objective was to evaluate clusterings of sizes 2-70, However, the stopping conditions limited the largest clusterings found for all samples to size 69 for iteration 1; 48 for iteration 2; and 41 for iteration 3. Figures S2-4, S2-5, and S2-6 display the distributions of normalized MED for the three iterations.

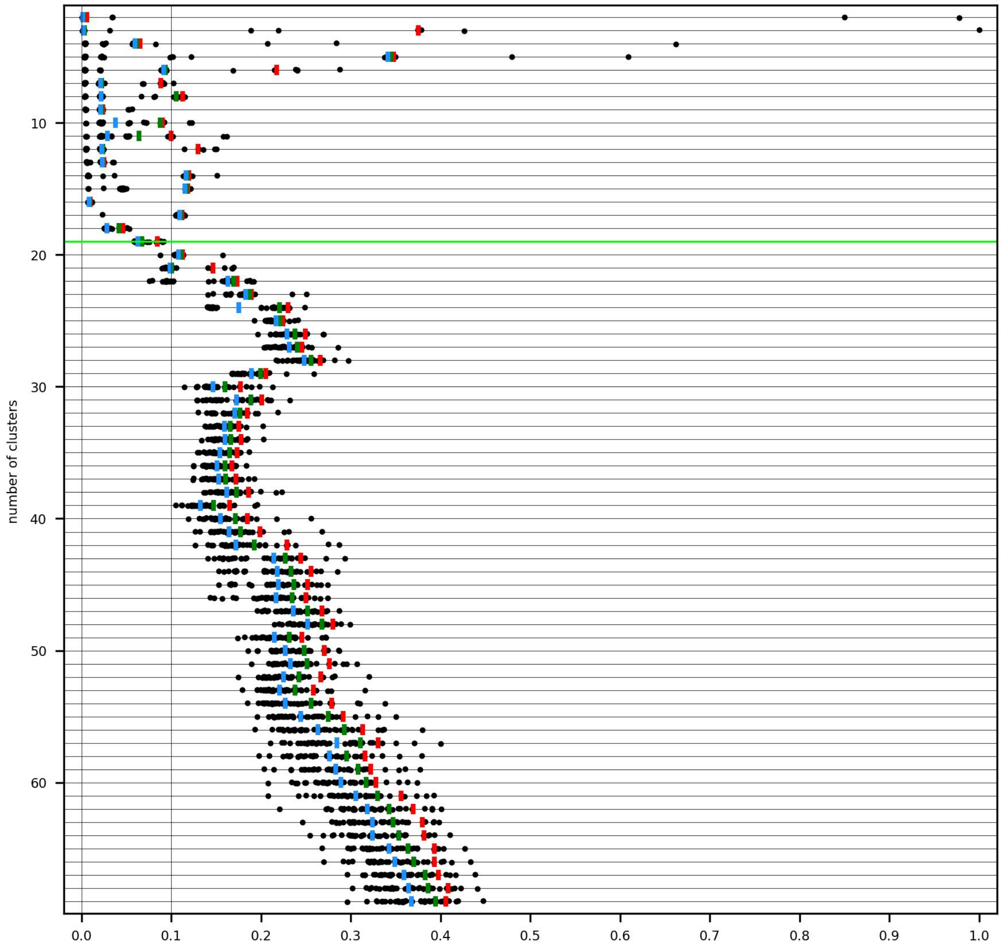


**Figure S2-4** 65k lung data (iteration 1): distributions of normalized MED for all generated clusterings; the largest stable clustering has size 19

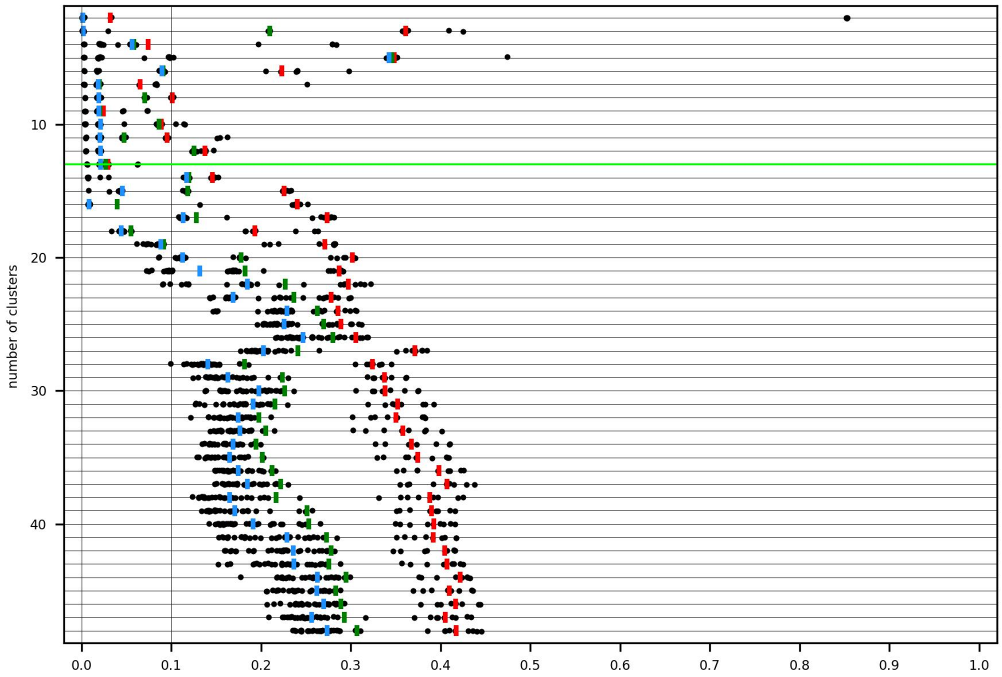


**Figure S2-5** 65k lung data (iteration 2): distributions of normalized MED for all generated clusterings; the largest stable clustering has size 13

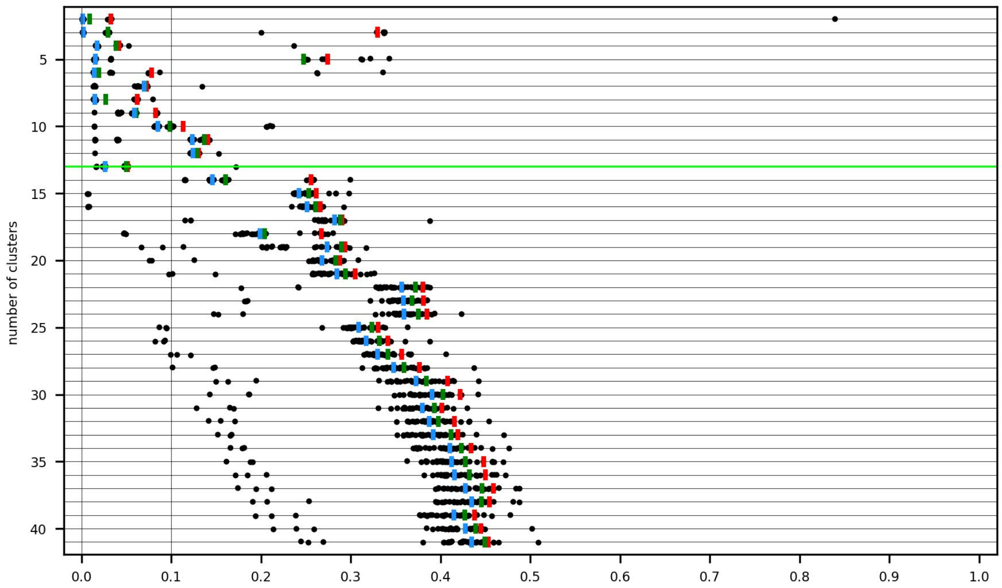


**Figure S2-6** 65k lung data (iteration 3): distributions of normalized MED for all generated clusterings; the largest stable clustering has size 13

**100k breast cancer**

As for the retinal and lung data, our objective was to evaluate clusterings of sizes 2-70, The stopping conditions limited the largest clusterings found for all samples to size 58 for iteration 1; 39 for iteration 2; and 51 for iteration 3. Figures S2-7, S2-8, and S2-9 display the distributions of normalized MED for the three iterations.

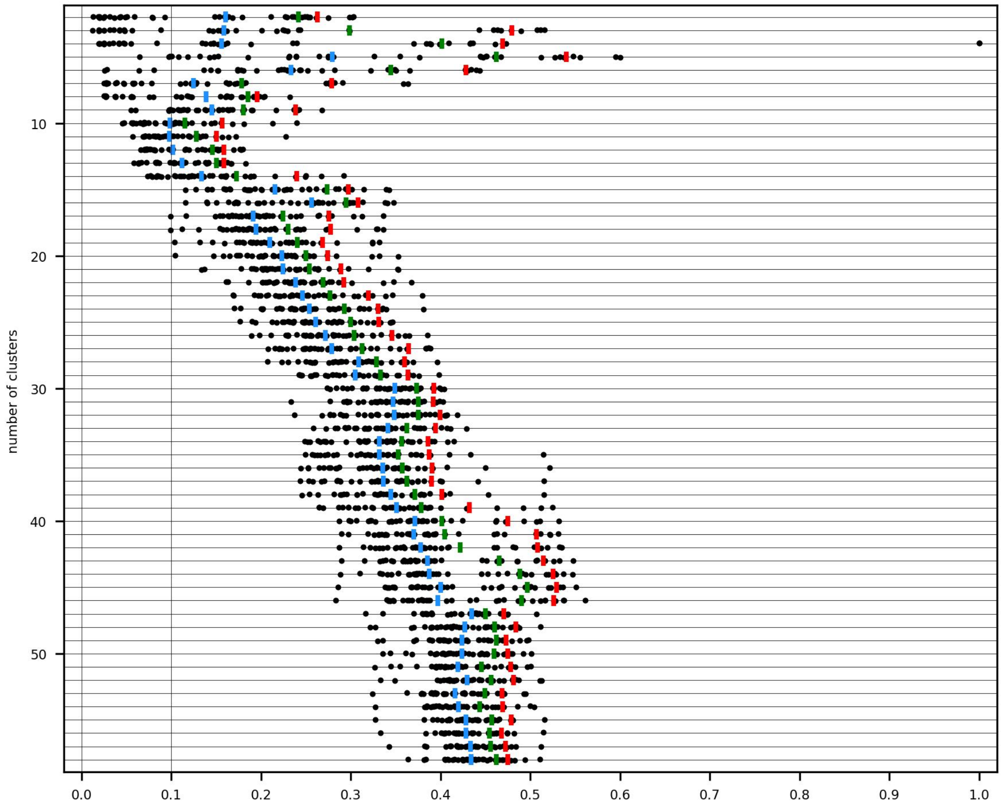


**Figure S2-7** 100k breast cancer data (iteration 1): distributions of normalized MED for all generated clusterings; the $90^{th}$ percentile (red) is greater than 0.10 for all all clusterings

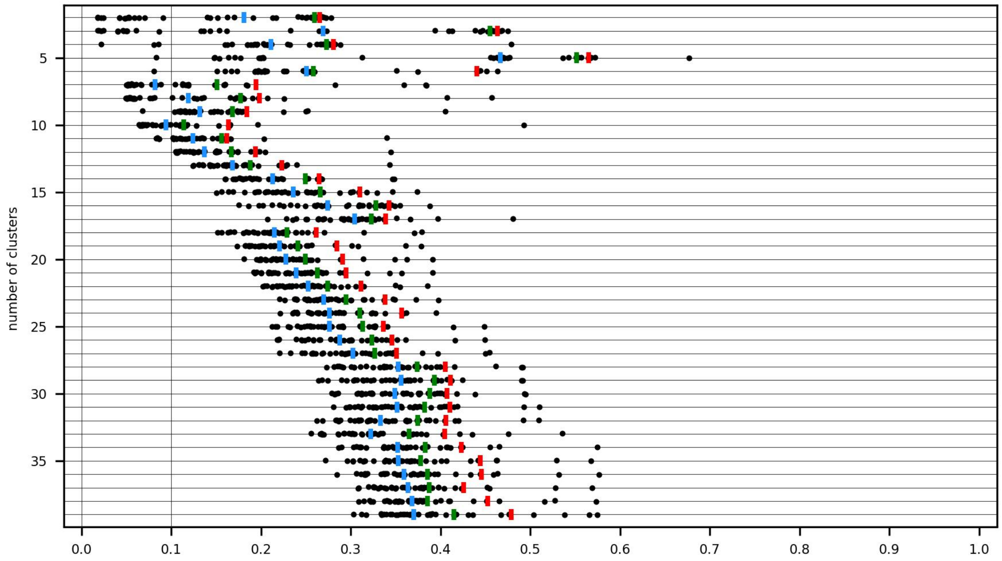


**Figure S2-8** 100k breast cancer data (iteration 2): distributions of normalized MED for all generated clusterings; the $90^{th}$ percentile (red) is greater than 0.10 for all all clusterings

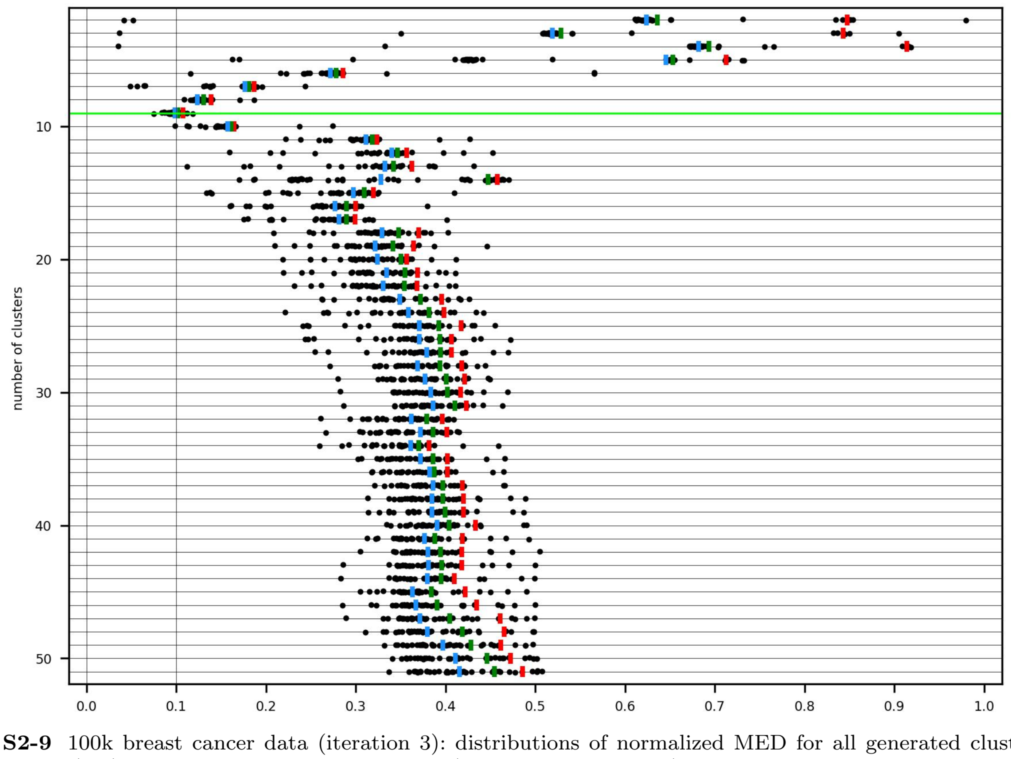


**Figure S2-9** 100k breast cancer data (iteration 3): distributions of normalized MED for all generated clusterings; the $90^{th}$ percentile (red) equals 0.107 for the 9-clustering (green highlighted line)
